# The curious case of a Chilean copepod (*Tigriopus* aff. *angulatus*) genome assembly

**DOI:** 10.64898/2026.03.11.711077

**Authors:** Isabelle P. Neylan, Rujuta V. Vaidya, Maheshi Dassanayake, Sergio A. Navarrete, Morgan W. Kelly, Brant C. Faircloth

## Abstract

*Tigriopus* copepods are found in splash pools on all seven continents from the equator to Arctic and Antarctic regions. Given their geographic distribution, frequent exposure to extreme environmental conditions, and strong signatures of local adaptation, these copepods have become models for exploring patterns of adaptation to stressful environments. However, most studies focus on a small subset of *Tigriopus* species, and there are few genome resources representing the diversity of *Tigriopus* species and populations. Here, we combine long-read, Pacific Biosciences HiFi data with short-read, Illumina HiC and RNA-seq data to assemble and annotate a genome representing a *Tigriopus* population from the coast of central Chile. Based on the level of divergence that we observed in mitochondrial genes, we also performed a comparison of morphological characteristics between this population and members of the *T. angulatus* complex. The assembly that we generated (qhTigAngs1.1.pri) includes 12 major scaffolds (N50 19Mbp, L50 7), equivalent to the number of chromosomes in other *Tigriopus* species. BUSCO and k-mer analyses of the assembly and BUSCO analyses of gene models are relatively complete (89-99%) with respect to gene or k-mer content. The level of divergence that we observed, morphological differences we recorded, and the spatial distance of this population from the type locality suggest that this Chilean population of *Tigriopus* may represent a novel species that we call *Tigriopus* aff. *angulatus*. These genomic resources will help us understand the diversity and structure of *Tigriopus* species and populations as well as facilitate future comparisons of adaptation across parallel environmental gradients.

## INTRODUCTION

Copepods are small marine or aquatic crustaceans and are some of the most abundant animals on the planet – making up a large part of marine food webs as dominant members of the zooplankton (Santhanam *et al*., 2019). Due to their pivotal role in marine ecosystem functioning, research often focuses on their ability to adapt and persist in the face of environmental stressors and climate change (DeMayo *et al*., 2021; Brennan *et al*., 2022). The genus *Tigriopus* is one group of Harpacticoid copepods that live in shallow splash pools in the high intertidal zone of rocky shores, and *Tigriopus* copepods (hereafter, *Tigriopus*) have become models for studying stress tolerance and ecophysiology because they are extremely tolerant of variation in temperature, salinity, and dissolved oxygen (Kelly *et al*., 2012; DeBiasse *et al*., 2018; Graham and Barreto, 2019) as well as to the presence of heavy metals (Raisuddin *et al*., 2007; Medina *et al*., 2008; Holan *et al*., 2016). *Tigriopus* are also an ideal system to explore the genetic mechanisms of local adaptation because they show strong inter-population differences in stress tolerance across latitudes, particularly in thermal tolerance (Schoville *et al*., 2012; Kelly *et al*., 2017; Griffiths *et al*., 2021). There is also significant population structure and a high potential for speciation in this genus because *Tigriopus* have extremely low gene flow (Kelly *et al*., 2013) and exhibit several molecular incompatibilities when hybrids are created across populations within a species (Barreto *et al*., 2018; Olsen *et al*., 2023).

Currently, there are 15 species of *Tigriopus* recognized (Walter & Boxshall, 2025) that have a global distribution spanning the Arctic to the Antarctic with members of the genus found on all seven continents (Nazari *et al*., 2021; Walter & Boxshall, 2025). Despite their importance to marine ecosystems and extraordinary distribution, studies of the phylogenetic relationships and global diversity of *Tigriopus* species and populations are few (Jung *et al*., 2006; Vecchioni *et al*., 2019), and only three species have reference genome assemblies publicly available: *T. californicus* (Barreto *et al*., 2018), *T. japonicus* (Jeong *et al*., 2020), and *T. kingsejongensis* (Kang *et al*., 2017). Most *Tigriopus* species are delineated by subtle morphological differences that are commonly described for individuals collected at one locality within a large range, despite observations throughout the clade of significant local adaptation (Schoville *et al*., 2012; Kelly *et al*., 2013; Vecchioni, 2024). Geography then serves as a proxy for these morphological differences during species identification (Wells, 2007), although the geographic range of certain species can encompass thousands of kilometers and include multiple continents.

The shortcomings of this approach are particularly evident among populations of *Tigriopus angulatus*, which is the binomial used to describe all *Tigriopus* found across southern oceans (Figure 1). Originally described on Macquarie Island and Tasmania by Lang (1933), the species has subsequently been identified on New Zealand’s South Island (Bradford, 1967), Marion Island off South Africa (Grindley, 1971), South Georgia Island off Argentina (Davenport *et al*., 1997), Kerguelen and Crozette Islands in the southern Indian Ocean (Soyer *et al*., 1987), King George Island off of the Antarctic Peninsula (Park *et al*., 2014), and purportedly in an Andean lake in Chile 3,000m above sea-level (Von Brehm, 1936). Subsequent work has split several populations of *T. angulatus* into new species, including *T. kerguelensis* and *T. crozettensis* in the southern Indian Ocean (Soyer *et al*., 1987), *T. raki* on the North Island of New Zealand (Bradford, 1967; Nazari *et al*., 2021), and *T. kingsejongensis* in Antarctica (Park *et al*., 2014). Given the huge geographic range of this group and high levels of local adaptation, it is likely that *T. angulatus* is a species complex (Soyer *et al*., 1987; Davenport *et al*., 1997; Park *et al*., 2014) that includes additional, distinct species.

**Figure 1.**
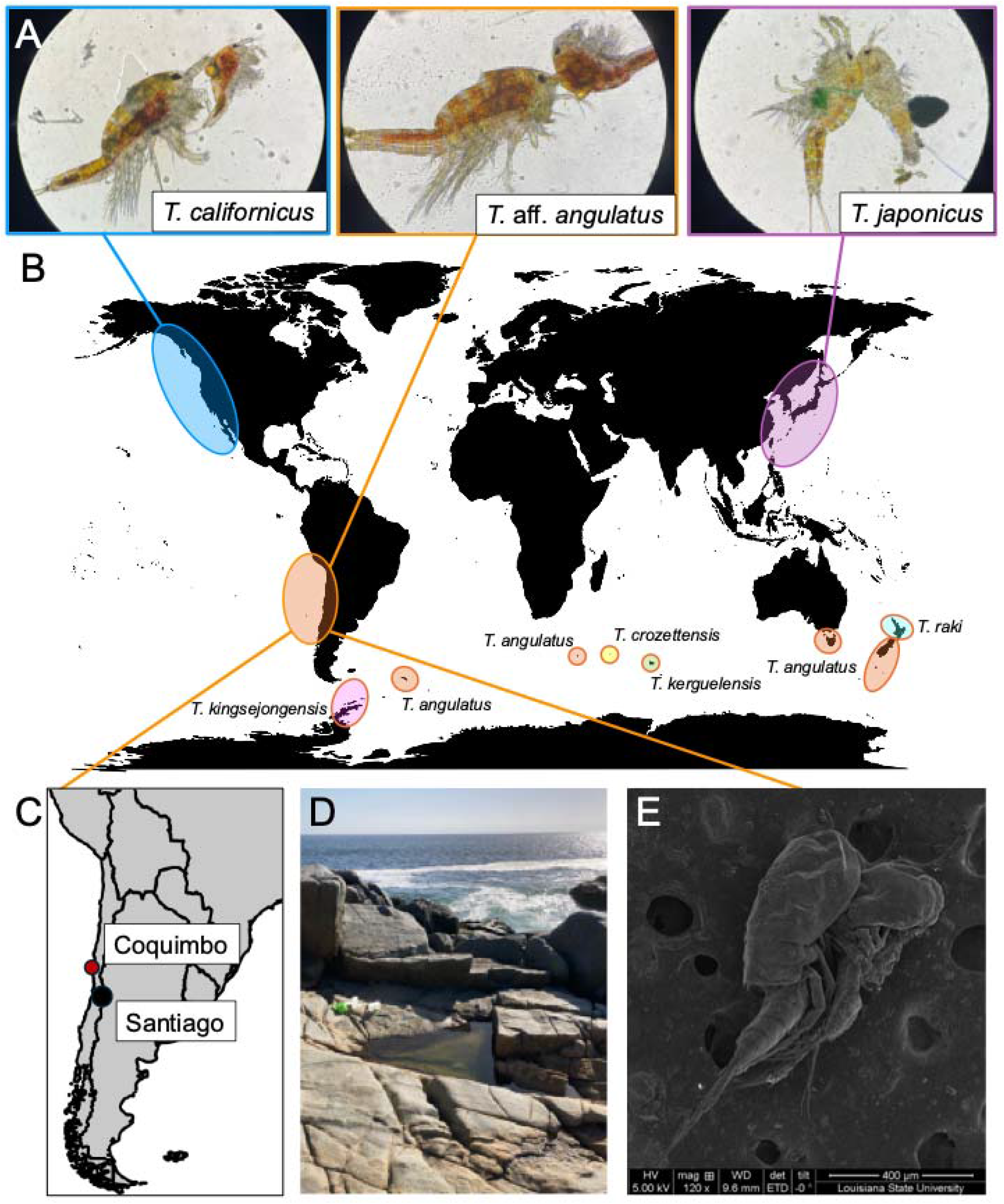
(A) Photos of male and female adult copepods taken with a compound microscope at 10X magnification of: *Tigriopus californicus* (in blue), Chilean *Tigriopus* that we designate *T.* aff. *angulatus* (in orange), and *T. japonicus* (in purple). (B) Collection localities of different populations of the *Tigriopus angulatus* species complex (orange circles) along with former members of the complex that have been named as unique species: *T. kingsejongensis* (pink shading), *T. crozettensis* (yellow), *T. kerguelensis* (green), and *T. raki* (teal). The inset map (C) and photo (D) show the location and habitat where we collected *T.* aff. *angulatus* individuals in Coquimbo, Chile. (E) A photo of a Chilean *T.* aff. *angulatus* mate-guarding pair captured using scanning electron microscopy.

Aside from the population described in the Andes, western South American *Tigriopus* are found in rocky splash pools along the Chilean coast (Figure 1) and have been observed from at least the Los Ríos region of south-central Chile (39.82°S, 73.17°W) to the regions of Arica and Parinacota in northern Chile (20.24°S, 70.13°W), although their populations likely extend further north into Peru (Fornes *et al*., 2010) and possibly Ecuador and Colombia. Given the absolute distance between these populations and the collection localities of the *T. angulatus* type specimens near New Zealand (>8000 km away), the degree of local adaptation observed within *Tigriopus* populations, and the identification of several new species within the *T. angulatus* complex (Soyer *et al*., 1987; Park *et al*., 2014; Nazari *et al*., 2021), *Tigriopus* from the western coast of South America likely represent one or several new species distinct from their counterparts inhabiting the high intertidal zones of other southern ocean coastlines.

Although relatively little is known about the phylogenetics, biology, and ecology of South American *Tigriopus*, previous work in Chile has explored their ability to withstand copper pollution (Medina *et al*., 2008) and measured lipid content across their ontogeny (Fornes *et al*., 2010). Outside of Chile, other studies using *T. angulatus* have demonstrated high heavy metal tolerance (Holan *et al*., 2016), low thermal tolerance compared to congeners (Davenport *et al*., 1997), and that adults can serve as an intermediate host for mosquito parasites (Wong & Pillai, 1980).

Our interest in South American *Tigriopus* populations derives from the fact that their latitudinal distribution (Fornes *et al*., 2010) mirrors that of their well-studied, North American congener, *T. californicus*, which has become a model for understanding thermal physiology, genetic adaptation, and mechanisms of adaptation and acclimation (Kelly *et al*., 2012, 2017; Schoville *et al*., 2012; Harada *et al*., 2019; Healy *et al*., 2019; Vaidya *et al*., 2025; Neylan *et al*., 2026). *Tigriopus* populations along the west coast of South America present a unique opportunity to explore how the mechanisms of adaptation and acclimation evolve in a naturally replicated thermal gradient (Navarrete *et al*., 2008; Tapia *et al*., 2014). This research could reveal important insights regarding parallel evolution, continue to advance our understanding of thermal adaptation mechanisms, and inform predictions for species persistence in the face of global change.

To enable these types of studies, we collected copepods representing a South American population of *Tigriopus* near Coquimbo, Chile and used these individuals to create an inbred line. We collected sequence data of various types from the inbred line and used these data to assemble and annotate a contiguous nuclear genome and mitochondrial genome representing this population. Because of the degree of mitochondrial divergence that we observed during the assembly process, we performed an analysis of mitochondrial sequences that included one former member of the *T. angulatus* species complex (*T. kingsejongensis*) as well as *T. californicus* and *T. japonicus*. Those results suggested substantial divergence between the Chilean *Tigriopus* and individuals representing *T. kingsejongensis* that mirrored the degree of divergence we observed among other *Tigriopus* species, so we used compound microscopy to diagnose key morphological differences between the Chilean individuals and other species within the *T. angulatus* complex. These combined analyses led us to conclude that the Chilean individuals likely represent a different species of *Tigriopus* that we refer to as *Tigriopus* aff. *angulatus*.

## METHODS

### Specimen collection and establishment of inbred lines

We collected copepods from splash pools near Coquimbo, Chile (30.067°S, 71.383°W; Figure 1C, D) during March of 2022 following methods described in Kelly et al. (2012). We brought the copepods back to the United States and maintained the population at Louisiana State University in an incubator (Percival 136VL) at 19° C under a 12hr light/12hr dark cycle using artificial seawater (Instant Ocean®) at a salinity of 35 ppt. To reduce the degree of heterozygosity in the individuals used for genome assembly, we created inbred lines from this Chilean population. Specifically, during August of 2023, we placed mate-guarding pairs into individual wells of a six-well plate (ThermoFisher Scientific™) to create six crosses. Once we detected the female was gravid, we removed males, and once offspring copepodites were visible, we removed the female to ensure sibling/sibling mating. We then moved three sibling/sibling mate-guarding pairs from each well to a new plate to form the next generation. We repeated this process for ten generations before placing the animals from each line into separate 950ml containers full of artificial seawater and maintaining these lines in an incubator at 19° C under a 12hr light/12hr dark cycle until sample collection.

### Nucleic acid extraction, library preparation, and sequencing

During February of 2024, we selected one line to use for DNA library preparation, and we sampled several replicates of 200 individuals from that line, which we placed into individual 1.5 µl microtubes. We subsequently flash-froze the replicate samples in liquid nitrogen and stored them at -80°C. We submitted one replicate to the Institute for Genome Sciences (IGS) at the University of Maryland School of Medicine where IGS staff extracted high-molecular weight DNA from the pooled individuals using a Qiagen MagAttract HMW DNA kit with a modified lysis procedure. Specifically, the pool of inbred individuals was thawed on ice, resuspended in 220 µl of Qiagen buffer ATL and 20 µL of proteinase K, vortexed, and mixed overnight on a thermomixer at 56°C and 900 RPM (approx. 17hrs total time). After incubation, specimens were chilled on ice, gently homogenized using a Dounce homogenizer, and centrifuged. Then, IGS staff transferred approximately 200 µL of supernatant to a clean 1.5mL Protein LoBind tube (Eppendorf AG) using a wide-bore 200 µL pipette tip. The remaining steps followed the MagAttract kit protocol for “Manual Purification of High-Molecular-Weight Genomic DNA from Fresh or Frozen Tissue”. Following extraction, IGS staff performed quality control on the extracted DNA and used the DNA to prepare a tagged SMRTBell library for HiFi sequencing without shearing. IGS sequenced the library on a partial Revio SMRT cell targeting 1-2 M reads, which we expected would yield ∼30-40X coverage assuming a genome size of ∼200-300 Mbp, which is similar to the genome size estimate of Tigriopus californicus (Barreto et al., 2018).

After we received the data from IGS, we removed adapter contamination from sequencing reads using cutadapt (Table 1; Martin, 2011), and we generated a temporary, collapsed assembly using hifiasm (Cheng *et al*., 2021, 2022). Then, we submitted a second replicate pool of individuals from the same inbred line to Phase Genomics (PG), where PG staff used the pooled samples to create a HiC library (Lieberman-Aiden *et al*., 2009; van Berkum *et al*., 2010) using the Phase Genomics Proximo Kit. After library preparation, PG staff generated a small number (< 1 M) of paired-end (PE) 150 bp sequencing reads from the library using an Illumina iSeq100 and aligned these reads to the temporary assembly to ensure the HiC library was of sufficient diversity for deeper sequencing. Following quality control, PG staff sent the library to a commercial provider who performed Illumina 150 base-pair, PE sequencing on a NovaSeq X Plus targeting 50 M read pairs. After receiving these data from PG, we trimmed sequence reads using cutadapt to remove the sequencing adapters, trailing bases having a quality score < 20, trailing ambiguous bases, and reads that were shorter than 40 bp.

**Table 1.**
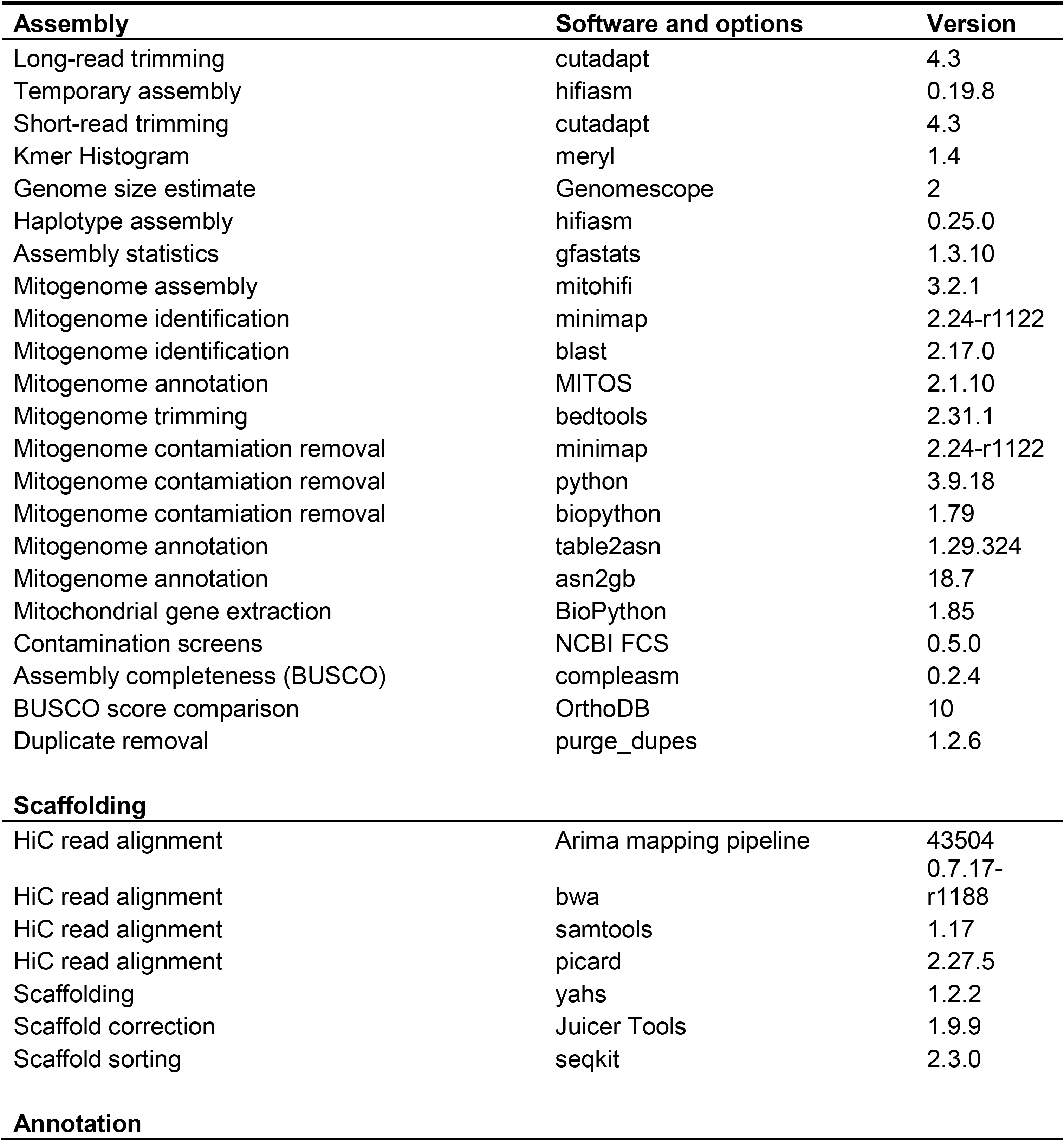
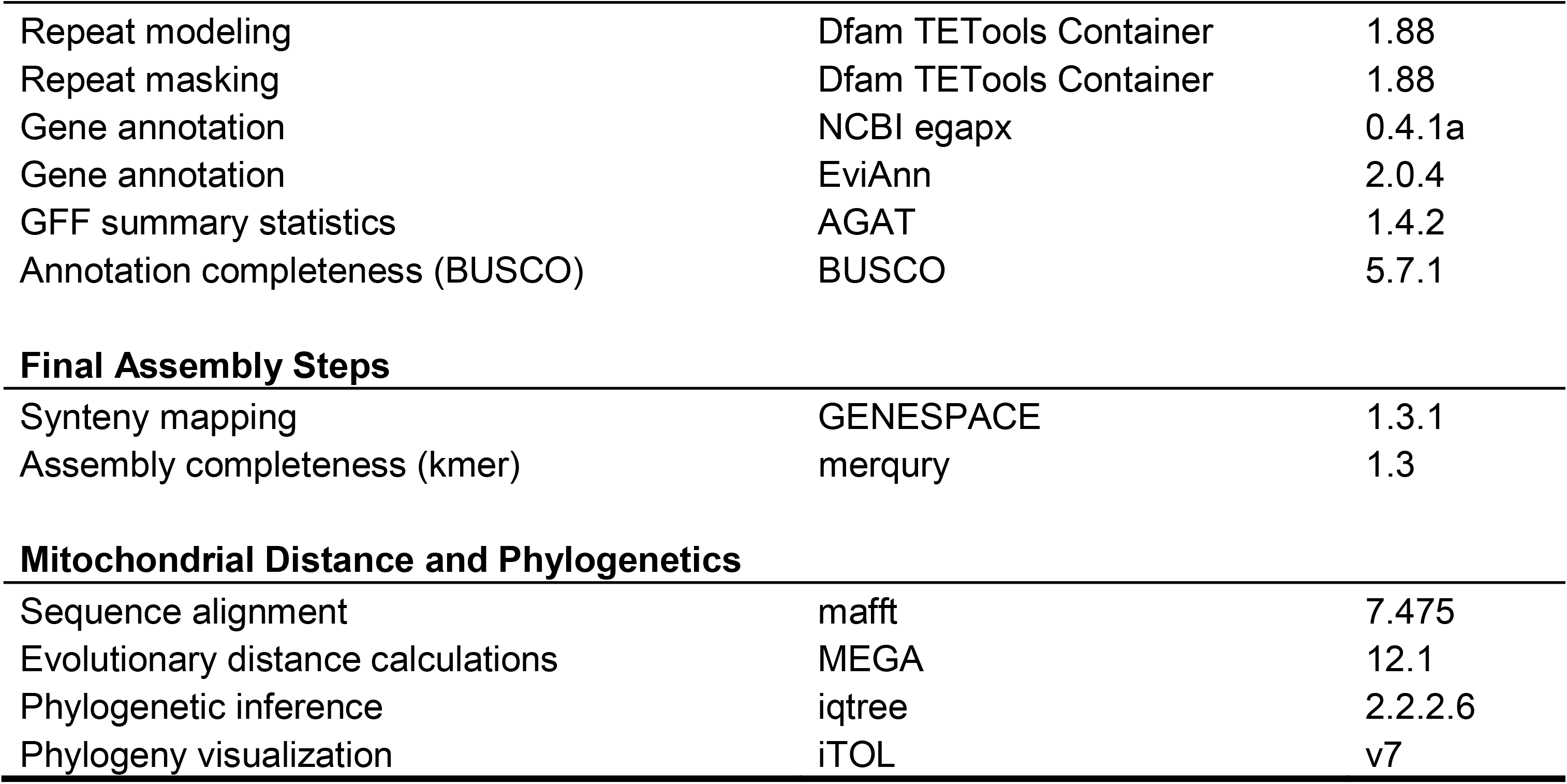
List of programs used to assemble, scaffold, annotate, and analyze components of the Chilean *Tigriopus* mitochondrial and nuclear genomes.

During May of 2025, we created six replicate pools by selecting 50-60 animals per pool from the same inbred line we used for genome sequencing, flash-freezing these samples in liquid nitrogen, and storing them at -80°C. We extracted RNA from replicates by homogenizing copepods using a TissueRuptor II (Qiagen, Inc.) followed by a protocol that combined steps 1-7 of the TRIzol protocol (Invitrogen, Inc.) with steps 4-11 of RNAeasy Plus kit protocol (Qiagen, Inc.; Neylan and Vaidya, 2025). We sent RNA extracts to Novogene Corporation Inc. (Sacramento, California), where Novogene staff analyzed RNA quality using a Bioanalyzer (Agilent, Inc.) with the Eukaryote Total RNA Nano chip. Novogene staff then prepared non-directional RNASeq libraries using poly-A tail selection from extracts that passed quality control, and they sequenced the resulting libraries on an Illumina NovaSeq X Plus, targeting 20 M, 150 base-pair, PE reads per library.

### Genome assembly

Prior to assembly, we estimated genome size and heterozygosity for the Chilean *Tigriopus* by generating a kmer histogram from the HiFi data using meryl (Rhie *et al*., 2020) with a kmer length of 21 and inputting the histogram to Genomescope (Ranallo-Benavidez *et al*., 2020). Then, we generated an assembly by inputting the HiFi and HiC data to hifiasm with default options. Although this procedure typically produces haplotype-phased assemblies, we were concerned that haplotype switching could occur due to the use of pooled DNA, so we performed all subsequent steps using the collapsed, primary assembly (“primary assembly” hereafter). After assembly, we used GNU awk to convert the graphical fragment assembly (GFA) file to FASTA format, and we computed assembly statistics using gfastats (Formenti *et al*., 2022).

We initially attempted to assemble the mitochondrial genome for the Chilean *Tigriopus* by inputting the trimmed HiFi reads to the containerized version of mitohifi (Uliano-Silva *et al*., 2023) with different reference genomes: *T. japonicus* (AB060648.1, Machida *et al*., 2002) *T. kingsejongensis* (MK598762.1, Hwang *et al*., 2019), and *T. californicus* (NC_008831.2, Burton *et al*., 2007). However, each attempt failed to identify sufficient reads aligning to the respective reference genome to enable assembly. Because hifiasm can assemble mitochondrial genomes from total DNA extracts as part of a genome assembly, we attempted to identify contigs of mitochondrial origin by aligning the temporary assembly to several mitochondrial reference assemblies (*T. japonicus, T. kingsejongensis*, and *T. californicus*) using minimap (Li, 2018) with the ‘-asm10’ preset. These searches did not return results, so we used standalone NCBI blastn (Camacho *et al*., 2009) with default parameters to search the temporary assembly for matches to the *T. japonicus* mitochondrial genome, which returned results for several contigs. We used faOneRecord from the Kent source (Kent *et al*., 2002) to extract the longest contig identified, and we uploaded the resulting FASTA to Galaxy (Abueg *et al*., 2024), where we annotated the contig using MITOS2 (Donath *et al*., 2019). The MITOS2 annotation in browser extensible data (BED) format suggested that small portions (100-200 bp) of the cytochrome c oxidase II (COX2) gene were duplicated several times at the start and the end of the linear contig that we extracted and annotated. So, we used bedtools (Quinlan and Hall, 2010) getfasta within Galaxy to trim these duplicated regions while retaining the single, annotated COX2 gene that was approximately the size we expected (∼700 bp) based on comparisons of COX2 sequences among the reference *Tigriopus* mitochondrial assemblies mentioned above. After trimming, we performed a second round of annotation with MITOS2 in Galaxy to ensure that annotation results were sensible, downloaded the resulting BED file, converted that to GenBank “feature table format” by hand, and used the NCBI table2asn and asn2gb programs to create a GenBank-formatted mitochondrial assembly file. The resulting mitochondrial assembly was somewhat shorter than we expected (14.1 kbp versus 14.5 kbp in *T. californicus* NC_008831.2; Burton *et al*., 2007), and it was hard to determine if it was circularized because the repetitive COX2 region that we trimmed was located at the overlapping ends of the contig, so we input this temporary reference to mitohifi along with the trimmed HiFi reads, and the mitochondrial assembly and annotation process completed successfully. We used BioPython (Cock *et al*., 2009; Van Rossum & Drake, 2011) to extract the cytochrome c oxidase I (COX1), cytochrome B (CYTB), and NADH dehydrogenase subunit 2 (ND2) sequences from the mitohifi assembly. Then, we input the COX1 sequence into the Barcode of Life Identification Engine (Ratnasingham *et al*., 2024) and all three gene sequences into the online version of NCBI blastn (Madden, 2003) to search for species-level matches and verify that the loci were similar to others for *Tigriopus*.

After assembling the mitochondrial genome, we used minimap to screen the primary assembly for mitochondrial contamination. We also screened the assembly for organismal and adapter contamination using the NCBI Foreign Contamination Screen (FCS) tools (Astashyn *et al*., 2024) with standard parameters, an NCBI taxon identification for *T. japonicus* (because *T. angulatus* was not entered in NCBI Taxonomy), and the NCBI GX database from 24 January 2023. We performed two rounds of screening using the NCBI FCS tool – during the first, we removed all hits marked by the software as “REVIEW” or “EXCLUDE” after ensuring that each were identified as prokaryotic in origin. During the second, we removed any remaining contigs that were identified as being prokaryotic in origin or that were identified as similar to *Tigriopus* food (i.e., flagged as algae).

After removing contaminants, we used gfastats to compute assembly statistics, and we ran compleasm (Huang and Li, 2023) to perform BUSCO (Simão *et al*., 2015) analyses using the arthopod BUSCOs from OrthoDBv10 (Manni *et al*., 2021). BUSCO results suggested that the duplication rate in the primary assembly was high, so we performed two successive rounds of duplicate purging using purge_dups (Guan *et al*., 2020), and we recomputed assembly statistics and BUSCO scores after each round to ensure that the two-step process had not been overly aggressive. Then, we used bwa (Li and Durbin, 2010), samtools (Danecek *et al*., 2021), and Picard (‘Picard toolkit’, 2019) within the Arima Genomics Mapping Pipeline (‘Arima-HiC Mapping Pipeline’, 2024) to align the trimmed, HiC data to the primary assembly, and we used YaHS (Zhou *et al*., 2023) and Juicer Tools (Durand *et al*., 2016) to scaffold the assembly and produce contact maps for the scaffolds. After ensuring that the contact map did not include scaffolds that appeared to be misassembled, we sorted the scaffolds by length from largest to smallest using seqkit (Shen *et al*., 2024). Scaffolding produced 12 large scaffolds (>14 Mbp) that were significantly larger (>6.5 times) than all remaining scaffolds and contigs. This number is equivalent to the number of chromosomes in other species of *Tigriopus* (Ar-rushdi, 1962; Foley *et al*., 2011), so based on these differences in size and the clear delineation of these scaffolds in the contact maps, we renamed each of these 12 scaffolds to reflect that they likely represent chromosomes. After renaming, we used Merqury (Rhie *et al*., 2020) to compute an estimate of k-mer completeness and quality, we recomputed assembly statistics and BUSCO scores using gfastats and compleasm, and we assigned a Tree of Life identifier to the assembly (Blaxter *et al*., 2024).

### Genome annotation

We used RepeatModeler (Flynn *et al*., 2020) within the Dfam Transposable Element Tools (TETools) container (‘TETools: Dfam Transposable Element Tools Docker container’, 2023) to model repeats in the primary assembly using the DFAM (Storer *et al*., 2021) and RepBase Repeat Masker libraries v20181026 (Bao *et al*., 2015). We input the resulting set of repeat models to RepeatMasker in the TETools container to create a general feature format (GFF) file of repeats and soft-mask repeats in the assembly. After repeat masking, we predicted gene models for the primary assembly by inputting the reads from the RNA-seq libraries and the NCBI taxonomy ID for *Tigriopus* (taxid: 6831) to the containerized version of the NCBI eukaryotic genome annotation pipeline (egapx). We also predicted gene models using the EviAnn software package (Zimin *et al*., 2025) with the option for functional annotation enabled, the binary alignment map (BAM) files for each RNA-seq library that were generated by NCBI egapx, and an evidence file that consisted of 896,885 protein sequences downloaded from the NCBI RefSeq database for taxonomic superclass multicrustacea (taxid: 2172821) on 20 October 2025. We compared summary statistics for protein coding genes between the GFF files output by NCBI egapx and EviAnn using the AGAT container (Dainat, 2022), and we compared BUSCO scores for the file of predicted proteins output by each program using a containerized version of BUSCO with the arthropod BUSCOs from OrthoDBv10.

### Genome comparisons

To enable comparisons between the Chilean *Tigriopus* assembly and those of other *Tigriopus* species that were available during June 2026, we downloaded the assemblies for *T. californicus* (GCF_007210705.1), *T. japonicus* (GCA_010645155.1), *T. kingsejongensis* (GCA_012959195.1), and *Tigriopus* “West” (GCA_051201535.1). Then, we computed assembly and completeness statistics for each using gfastats and the BUSCO container with the Arthropoda OrthoDBv10 BUSCOs. We also generated a figure illustrating the degree of synteny in gene order between the assembly representing the Chilean *Tigriopus* and *T. californicus* using GENESPACE (Lovell *et al*., 2022). We were unable to include other assemblies in this analysis because none were annotated. Other, relatively simple methods of generating synteny plots using alignments of nucleotide sequences failed to produce reasonable alignments between assemblies at levels of sequence divergence ≤5%.

### Mitochondrial distance and phylogeny

Given how difficult it was to assemble the mitochondrial genome, we were interested in estimating the pairwise evolutionary distances between the Chilean *Tigriopus* and the reference *Tigriopus* mitochondrial assemblies mentioned above. So, we extracted COX1 from each mitochondrial assembly using BioPython (Chilean *Tigriopus*) or the online tools at NCBI GenBank (other assemblies); aligned the sequences using mafft (Katoh and Standley, 2013) with the ‘--adjustdirection’, ‘--maxiterate 1000’, ‘--localpair’ options; and used MEGA (Stecher *et al*., 2025) with the invertebrate mitochondrial code to estimate the pairwise distance between them by calculating the proportion of nucleotide sites at which they were different (p-distance) or using the Jukes-Cantor model (Jukes and Cantor, 1969).

We were also interested in inferring the phylogenetic relationships among *Tigriopus* species, so we took two approaches. First, we wanted to infer relationships using many nuclear loci, so we extracted Arthropoda OrthoDBv10 BUSCOs from one outgroup individual (*Acartia tonsa*, GCF_053477335.1) using the BUSCO container; combined these results with the BUSCOs we extracted from other *Tigriopus* genomes; and input the results to the BUSCO phylogenomics tool (https://github.com/jamiemcg/BUSCO_phylogenomics) to identify, align, and concatenate BUSCO protein sequences that were present in at least four of six (65%) assemblies. Then, we used IQ-TREE (Nguyen *et al*., 2015) to identify the best site-rate substitution model given the protein data (-m MFP), infer the best maximum likelihood phylogeny, generate 1000 ultrafast bootstrap replicates, and reconcile the best phylogeny with the bootstrap replicates. The BUSCO phylogenomics tool also produces gene trees for extracted BUSCOs using fasttree, and we input these gene trees to Astral-IV+Castles-2 (Zhang *et al*., 2025) to infer a coalescent phylogeny with branch lengths and local branch support. We renamed tips in the resulting phylogenies with the taxon to which they were assigned in GenBank, and we rooted both trees on the outgroup taxon.

For the second approach, we wanted to infer a phylogeny using a larger sample of individuals, so we used searches of NCBI GenBank to download COX1 sequences for all *Tigriopus* individuals that appeared to be unique as of January 2026, as well as COX1 sequences for *Acartia tonsa*, *Caligus clemensi*, and *Lepeophtheirus salmonis*, which we used as outgroup individuals (Edmands, 2001; Machida *et al*., 2002; Jung *et al*., 2006; McBeath *et al*., 2006; Burton *et al*., 2007; Denis *et al*., 2009; Handschumacher *et al*., 2010; Peterson *et al*., 2013; Park *et al*., 2014; Baek *et al*., 2016; Karanovic *et al*., 2018; Hwang *et al*., 2019; Vecchioni *et al*., 2019; Figueroa *et al*., 2020). We aligned the COX1 sequences with mafft using the same options that we used to estimate pairwise distance, and we used IQ-TREE to identify duplicated sequences in the alignment. After retaining only unique sequences, we ran IQ-TREE again, with the options to select the best fitting site-rate substitution model given the nucleotide data (-m MFP), infer the best maximum likelihood phylogeny, generate 100 standard bootstrap replicates (Felsenstein, 1985), and reconcile the best phylogeny with the bootstrap replicates. We renamed tips in the resulting phylogeny with the taxon to which they were assigned in GenBank, their GenBank accession number, and their rough collection locality – based either on GenBank metadata or the publication for which they were collected. We rooted the tree on the branch uniting the outgroup taxa, and we used the Interactive Tree of Life web application (Letunic and Bork, 2024) to collapse branches having bootstrap support < 70% (Hillis and Bull, 1993).

### Morphological comparisons

Again, because of the difficulty we had assembling the mitochondrial genome and also because of the differences we observed after estimating evolutionary distances and inferring phylogenies, we were interested in comparing key morphological traits between individuals representing the Chilean *Tigriopus* population and other *Tigriopus* species. Morphological data are the primary mechanism used to differentiate taxa within the *T. angulatus* complex and currently serve as the only way to make reasonable comparisons among them. Therefore, we mounted multiple male and female adult individuals representing the population of Chilean *Tigriopus* from our lab cultures onto microscope slides for examination under a compound microscope and used dichotomous keys and morphological descriptions to collect data regarding key morphological traits. Then, we compared these measurements with the same trait data from other *Tigriopus* species within the *T. angulatus* species complex that we identified from the literature (Soyer *et al*., 1987; Davenport *et al*., 1997; Wells, 2007; Chullasorn *et al*., 2012; Park *et al*., 2014; Nazari *et al*., 2021) as well as *T. californicus* and *T. japonicus* individuals from cultures that we maintain in the laboratory. These laboratory cultures were derived from *T. californicus* collected from Bird Rock near San Diego, California USA (32.815°N, 117.273°W) during summer of 2022 and *T. japonicus* collected from Hong Kong (22.339°N, 114.267°E) during summer of 2019.

## RESULTS

### Sequencing, assembly, and annotation

We collected a total of 1.3 M HiFi sequencing reads from the inbred line representing the Chilean *Tigriopus* that had mean length of 9,154 (SD ± 4295) and totaled 12.0 M base pairs. Cutadapt did not identify HiFi reads containing adapters. Illumina sequencing of the HiC library produced 52.3 M PE150 read pairs, and Illumina sequencing across six RNA-seq libraries representing different pools of individuals produced a total of 136.1 M Illumina PE150 reads (mean per library: 22.7 M; SD ± 1.2 M).

Genomescope estimated that the maximum haploid length of the Chilean *Tigriopus* genome was 246.6 Mbp and that heterozygosity of the DNA used for sequencing was 0.46-0.48%. This genome size estimate suggested the HiFi sequencing coverage was approximately 49X. The length of the primary contig assembly was larger than this estimate (358 Mbp; Supplemental Table 1). The mitochondrial genome assembly for the Chilean *Tigriopus* was 16,534 bp in length, contained 36 genes, and was identified by mitohifi as being circular. The Barcode of Life Identification Engine did not identify matches to the COX1 sequence we extracted, but the top blastn hits for the three mitochondrial loci we extracted (COX1, CYTB, ND2) were other *Tigriopus* species. The NCBI adapter screen did not identify adapters in any contigs, but the NCBI organismal contamination screen identified many contaminating contigs. Most of the contaminating contigs corresponded to bacteria (*Leptolyngbya* sp., *Tateyamaria* sp.), which we expected because we did not treat inbred lines with antibiotics prior to creating the pool of individuals for DNA extraction. We removed 228 contaminating contigs (30.8 Mbp) from the primary assembly during the first round of FCS cleaning and 45 contaminating contigs (3.2 Mbp) during the second round.

After contaminant removal, the size of the primary assembly remained larger than the genome size estimate, and BUSCO analyses suggested that this difference resulted from the larger number of duplicate contigs (Supplemental Table 1). The elevated rate of contig duplication we observed was likely because we extracted DNA from a pool of several hundred inbred individuals to yield sufficient DNA for library preparation. The first round of duplicate purging removed 388 contigs (70.7 Mbp) from the primary assembly, and the second round removed 17 contigs (6.7 Mbp)., After purging, the total length for the assembly was very close (± 1%) to the Genomescope predictions and substantially improved the BUSCO score by reducing the number of single-copy duplicates (Supplemental Table 1). The scaffolding procedure increased contiguity of the purged, primary assembly (Scaffold N50 ∼ 19 Mbp; Supplemental Table 1), and we identified 12 large scaffolds (>14 Mbp; Supplemental Figure 1), which is equivalent to the number of chromosomes in other species of *Tigriopus* (Ar-rushdi, 1962; Foley *et al*., 2011). After naming these 12 major scaffolds as chromosomes, based on their size, we assigned a versioned Tree of Life identifier to the final assembly: qhTigAngs1.1.pri. The final set of summary statistics BUSCO scores, k-mer completeness estimates, and quality values (QVs) suggested qhTigAngs1.1.pri was contiguous and reasonably complete (Table 2).

**Table 2.**
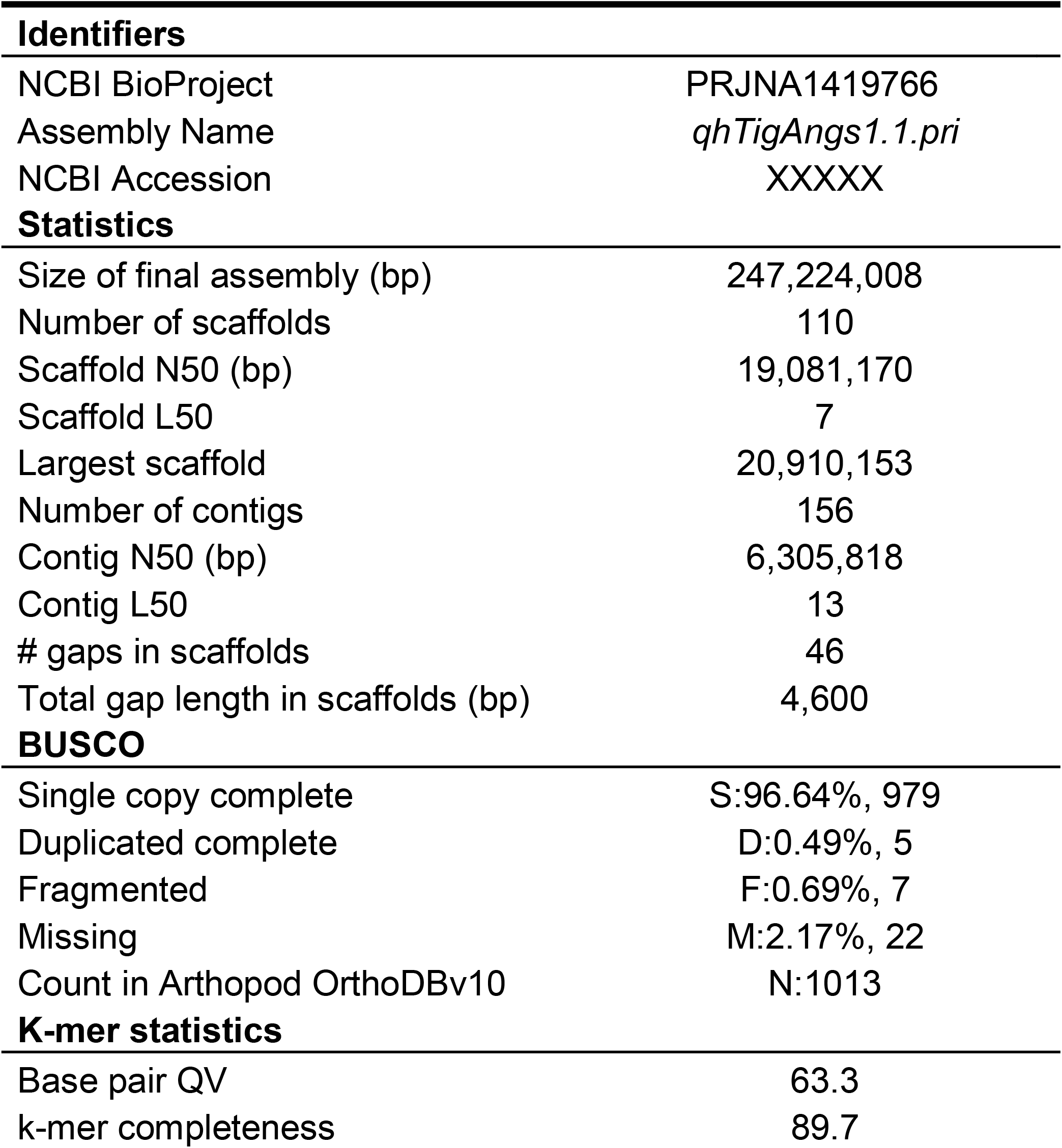
Summary of NCBI identifiers and assembly statistics including completeness and quality metrics for both haplotypes. Additional details are provided in Supplemental Table 1.

| **Identifiers** |  |
| --- | --- |
| NCBI BioProject | PRJNA1419766 |
| Assembly Name | *qhTigAngs1.1.pri* |
| NCBI Accession | XXXXXX |
| **Statistics** |  |
| Size of final assembly (bp) | 247,224,008 |
| Number of scaffolds | 110 |
| Scaffold N50 (bp) | 19,081,170 |
| Scaffold L50 | 7 |
| Largest scaffold | 20,910,153 |
| Number of contigs | 156 |
| Contig N50 (bp) | 6,305,818 |
| Contig L50 | 13 |
| # gaps in scaffolds | 46 |
| Total gap length in scaffolds (bp) | 4,600 |
| **BUSCO** |  |
| Single copy complete | S:96.64%, 979 |
| Duplicated complete | D:0.49%, 5 |
| Fragmented | F:0.69%, 7 |
| Missing | M:2.17%, 22 |
| Count in Arthropod OrthoDBv10 | N:1013 |
| **K-mer statistics** |  |
| Base pair QV | 63.3 |
| k-mer completeness | 89.7 |

Repeat modeling and mapping identified 43% of qhTigAngs1.1.pri as repeats (Supplementary Table 2), most of which were LINEs (14%) or LTR elements (14%). The NCBI egapx program predicted 14,839 protein coding gene models for qhTigAngs1.1.pri while EviAnn predicted 14,261 models, and BUSCO scores for each set of predicted proteins suggested that both were relatively complete (95-96% complete BUSCOs; Supplementary Table 3).

Comparisons among *Tigriopus* assemblies (Supplemental Table 4) showed that qhTigAngs1.1.pri is relatively contiguous, particularly with respect to the contig N50s. When comparing synteny between qhTigAngs1.1.pri and the assembly for *T. californicus* (Supplemental Figure 2), there also appear to be few chromosomal translocations. Interestingly, inversions within certain chromosomes seem frequent (e.g., 1-7, 9), while other chromosomes (e.g., 8, 12) appear to maintain identical gene order between the two species.

### Mitochondrial distance and phylogeny

The estimates of evolutionary distance that we computed (Supplemental Table 5-6) indicated substantial differences among the COX1 locus extracted from the Chilean *Tigriopus* mitochondrial assembly relative to COX1 loci representing *T. japonicus, T. kingsejongensis*, and *T. californicus*. These results suggest that the Chilean *Tigriopus* population we sampled is most closely related to *T. kingsejongensis*, although the evolutionary distances between these two taxa were substantial and comparable to the distances we observed between other pairs of different *Tigriopus* species.

We extracted 979 BUSCO loci from five *Tigriopus* genomes and the outgroup genome (*Acartia tonsa*) where each locus was present in four of six taxa. We aligned these to produce a character matrix that included 66,359 parsimony informative sites. Model selection identified the Q.INSECT+F+I+R4 site-rate substitution model as the one that best fit the data. Concatenated and coalescent phylogenies (Supplemental Figure 3) suggested that the Chilean *Tigriopus* population we sampled is sister to *Tigriopus kingsejongensis*. Branch lengths inferred using both methods also suggested divergence between the Chilean *Tigriopus* and *T. kingsejongensis* is similar to the levels of divergence we observed between other *Tigriopus* species (e.g., *T. californicus* versus *T. japonicus*).

We identified a total of 154 unique COX1 sequences collected from at least five different *Tigriopus* species (Supplemental Table 7), and we aligned these to produce a character matrix that included 662 parsimony informative sites. Model selection identified the TIM+F+I+G4 site-rate substitution model as the one that best fit the data. The resulting phylogeny using this model (Figure 2; Supplemental Figure 4) suggested that the Chilean *Tigriopus* population we sampled is sister to, but not nested within, a clade of *Tigriopus kingsejongensis*. Similar to our estimates of evolutionary distance, the branch lengths for these two taxa illustrate substantial divergence at the COX1 gene that are comparable to or exceed the differences between other *Tigriopus* species.

**Figure 2.**
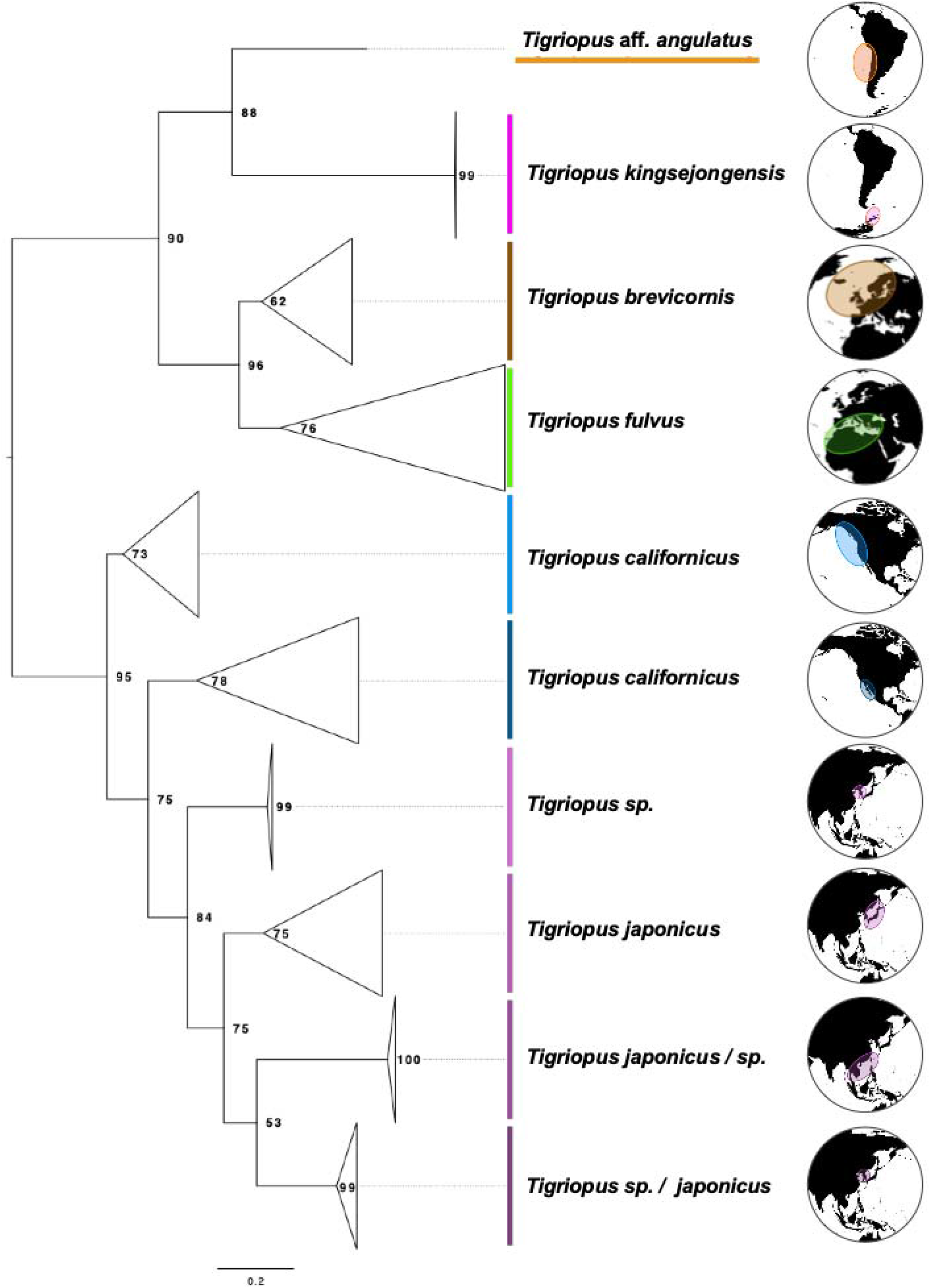
A phylogeny of *Tigriopus* inferred from unique COX1 sequences identified in NCBI GenBank and the COX1 sequence we extracted from the mitochondrial assembly of Chilean *Tigriopus* that we designate *T.* aff. *angulatus*. Clades have been collapsed to represent different species or different populations, and the map images to the right of taxon names shows approximate collection localities. We retained the specific epithet assigned to sequences by GenBank submitters, and when a specific epithet appears along with “sp.”, the order shows the taxon to which most species in the clade were assigned, with the left-most term representing the majority. Values at the nodes show bootstrap support, and a version of the phylogeny that has not been collapsed into major clades is available as Supplemental Figure 4.

### Morphological comparisons

As expected, the data that we collected for key morphological traits suggested the Chilean *Tigriopus* population was more similar to other species and populations within the *T. angulatus* complex than to *T. californicus* or *T. japonicus* (Figure 3). For example, there are substantial differences between taxa in the *T. angulatus* complex and *T. californicus* or *T. japonicus* in the number of: (1) setae on the endopod of the mandible (e.g., 2 inner setae for *T. angulatus* versus 3 for *T. californicus and japonicus*) and (2) setae on P4 exopod segment 3 (a total of 8 versus 7). However, there are also morphological differences within the *T. angulatus* complex, with the Chilean *Tigriopus* differing from the austral *T. angulatus* type in two key traits: (1) the number of setae on the exopod of their antenna (A2; 1:1:3 on segments one, two, and three for the *T. angulatus* type versus a pattern of 2:1:2 for the Chilean *Tigriopus*) and (2) the absence of a well-developed knob on the endopod of P2 in males. Historically, these differences have been sufficient to describe new species that were formerly described as populations of *T. angulatus* (Soyer *et al*., 1987; Park *et al*., 2014). And, among all the taxa that currently or formerly comprise *T. angulatus*, Chilean *Tigriopus* appear to possess a distinct combination of key diagnostic traits.

**Figure 3.**
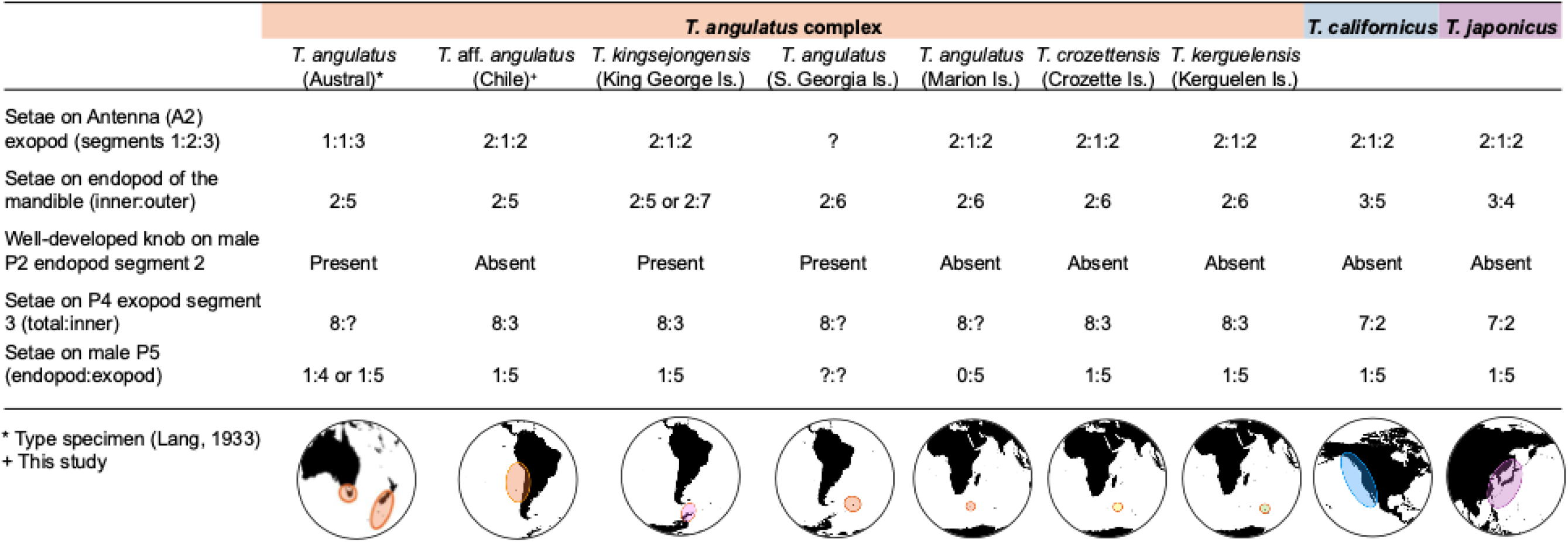
Morphological comparisons of key diagnostic traits for populations and species currently or previously within the *Tigriopus angulatus* complex as well as *T. californicus* and *T. japonicus.* We collected diagnostic trait data using a compound microscope (10-40X magnification) from Chilean *Tigriopus* (*T.* aff. *angulatus*), *T. californicus*, and *T. japonicus* and compared these observations to descriptions of the same traits in other species and populations of the *T. angulatus* complex that we identified in the literature (see Methods section for references). Measurements that we could not find in the literature are indicated by a “?”. The map images at the bottom of the figure show approximate collection localities for each species or population.

## DISCUSSION

We were initially interested in assembling a genome that represented a Chilean *Tigriopus* population to study how these organisms adapt to changing environments and how the molecular mechanisms underlying these adaptations evolve, as well as how these mechanisms may be similar to or different from those that enable *T. californicus* populations to survive along a comparable thermal gradient spanning the western coast of North America. The assembly that we generated should facilitate these types of studies because it is contiguous and relatively complete (Table 2; Supplemental Table 1); includes 12 large scaffolds (>14 Mbp) equivalent to the number of chromosomes observed in other species of *Tigriopus* (Barreto *et al*., 2018); and has reasonable sets of relatively complete gene models (Supplemental Table 3).

Perhaps more interesting than the nuclear genome assembly were the difficulties we encountered trying to assemble a mitochondrial genome for this taxon and our subsequent observations of significant mitochondrial divergence (Figure 2; Supplemental Tables 5-6) between Chilean *Tigriopus*, *T. kingsejongensis*, and other *Tigriopus* species. While the extent of mitochondrial divergence we observed was unexpected, divergence is typically higher in *Tigriopus* than in most other taxa (Barreto *et al*., 2018). For example, across populations of *T. californicus*, previous work has reported >20% nucleotide divergence and >25% amino acid substitutions in mitochondrial DNA sequences (Burton *et al*., 2007) as well as rates of mitochondrial substitution that far exceed the rates of nuclear substitution (Willett, 2012). These mitochondrial mutation rates are linked to multiple mito-nuclear incompatibilities across *T. californicus* populations driving reproductive isolation and likely contributing to strong patterns of local adaptation (Lima *et al*., 2019).

Unfortunately, mitochondrial and nuclear genomes representing the diversity of *Tigriopus* species and populations are few, and this limits our understanding of how rates of molecular evolution differ among these groups as well as the evolutionary relationships between *Tigriopus* species and populations – which also confuses their taxonomy. Because it included multiple individuals representing *T. kingsejongensis*, our mitochondrial analysis (Figure 2; Supplemental Figure 4) suggests that Chilean *Tigriopus* are sister to (and not nested within) *T. kingsejongensis*, which are the closest (∼3,700 km) *Tigriopus* species for which genetic data exist. The levels of mitochondrial and nuclear sequence divergence that we observed between the Chilean *Tigriopus* and *T. kingsejongensis* are similar to those observed between other *Tigriopus* species and suggest that Chilean *Tigriopus* and *T. kingsejongensis* represent different species. Because of their location in the Southern Ocean, prior to their description *T. kingsejongensis* would have been considered a population of *T. angulatus* (Park *et al*., 2014). Phylogenies we (Figure 2; Supplemental Figure 1) and others have inferred clarify some aspects of the Tigriopus relationships, but the available data limit a clear understanding of what is likely a complex set of relationships among current or former members of the *T. angulatus* complex. Collection of genome-scale data across the global distribution of *Tigriopus* would help to alleviate these shortcomings.

The degree of mitochondrial divergence we observed (Figure 2; Supplemental Table 5-6) and the lack of molecular data from taxa currently or formerly referred to as *T. angulatus* encouraged us to examine a set of key morphological traits that are typically used to diagnose *Tigriopus* species (Chullasorn *et al*., 2012; Park *et al*., 2014; Nazari *et al*., 2021). These characters suggest that the Chilean *Tigriopus* are unique when compared to *T. angulatus* and species formerly within the *T. angulatus* complex (Figure 3), and the differences we observed between Chilean *Tigriopus* and *T. angulatus* are similar in degree to those used to name new *Tigriopus* species.

Given these morphological differences, combined with the geographical distance between the collection localities of the *T. angulatus* type specimen and the Chilean *Tigriopus* individuals, we are reasonably confident that the Chilean *Tigriopus* population we sampled represents a distinct species. However, because there are no molecular data representing *T. angulatus* from the type locality (Macquerie Island; Lang, 1933) and because there are almost no molecular data for most remaining current and former members of the *T. angulatus* species complex (cf. *T. kingsejongensis*), it does not seem appropriate to name a new species at this time. As a result, we refer to this Chilean population as *Tigriopus* aff. *angulatus*.

Taken together, our results suggest that work is needed to better understand the global phylogeny of *Tigriopus*, particularly the relationships among *Tigriopus* populations spanning the southern oceans. At a more local scale, our results also suggest that future studies of *Tigriopus* aff. *angulatus* along the Chilean coast should delineate the extent of this species, including whether *Tigriopus* populations inhabiting the remainder of the western South American coastline are the same or different species. Finally, we hope that the genome assembly and annotation resources we generated will spur the development of *Tigriopus* aff. *angulatus* into a tractable model for studying whether parallel changes underlie adaptations to changing environments and ultimately contribute to our understanding of species persistence in a changing world.

## FUNDING

This work was supported by National Science Foundation award IOS-2154283 to MWK, MD, and BCF as well as a National Science Foundation Postdoctoral Research Fellowship in Biology 2305966 to IPN. Portions of this research were conducted with high-performance computational resources provided by Louisiana State University (http://www.hpc.lsu.edu).

## ACKNOWLEDGEMENTS

We thank Xuechu (Ellie) Zhao, Lisa Sadzewicz, and Luke Tallon at the IGS for their help collecting long-read sequencing data and Anna Schultz, Hayden Davis, and Kayla Young at Phase Genomics who helped collect the short-read HiC data. We also thank Emma Crable for assistance preparing the inbred lines and maintaining copepod cultures in the lab. Finally, we thank two anonymous referees and Associate Editor Kimberly Andrews for their comments that improved this manuscript.

## DATA AVAILABILITY

Sequence data are available from the NCBI as part of BioProject PRJNA1419766. The assembly has been deposited at DDBJ/ENA/GenBank as part of BioProject PRJNA1423750 and under the accessions XXXXXXXX. The version described in this paper is QQQQ. The supplemental tables, a list of steps used to assemble and annotate the genome, RepeatMasker annotations, gene models, and phylogenies in Newick format are available from FigShare (https://doi.org/10.6084/m9.figshare.29217905).

## Notes

https://doi.org/10.6084/m9.figshare.29217905

### Competing Interest Statement

The authors have declared no competing interest.

### Summary of Updates

We now report a primary (collapsed) assembly rather than two, haplotype-phased assemblies. We have added three new supplemental figures.

