## Supplemental Figure; Supplemental Table for "The curious case of a Chilean copepod (*Tigriopus* aff. *angulatus*) genome assembly"

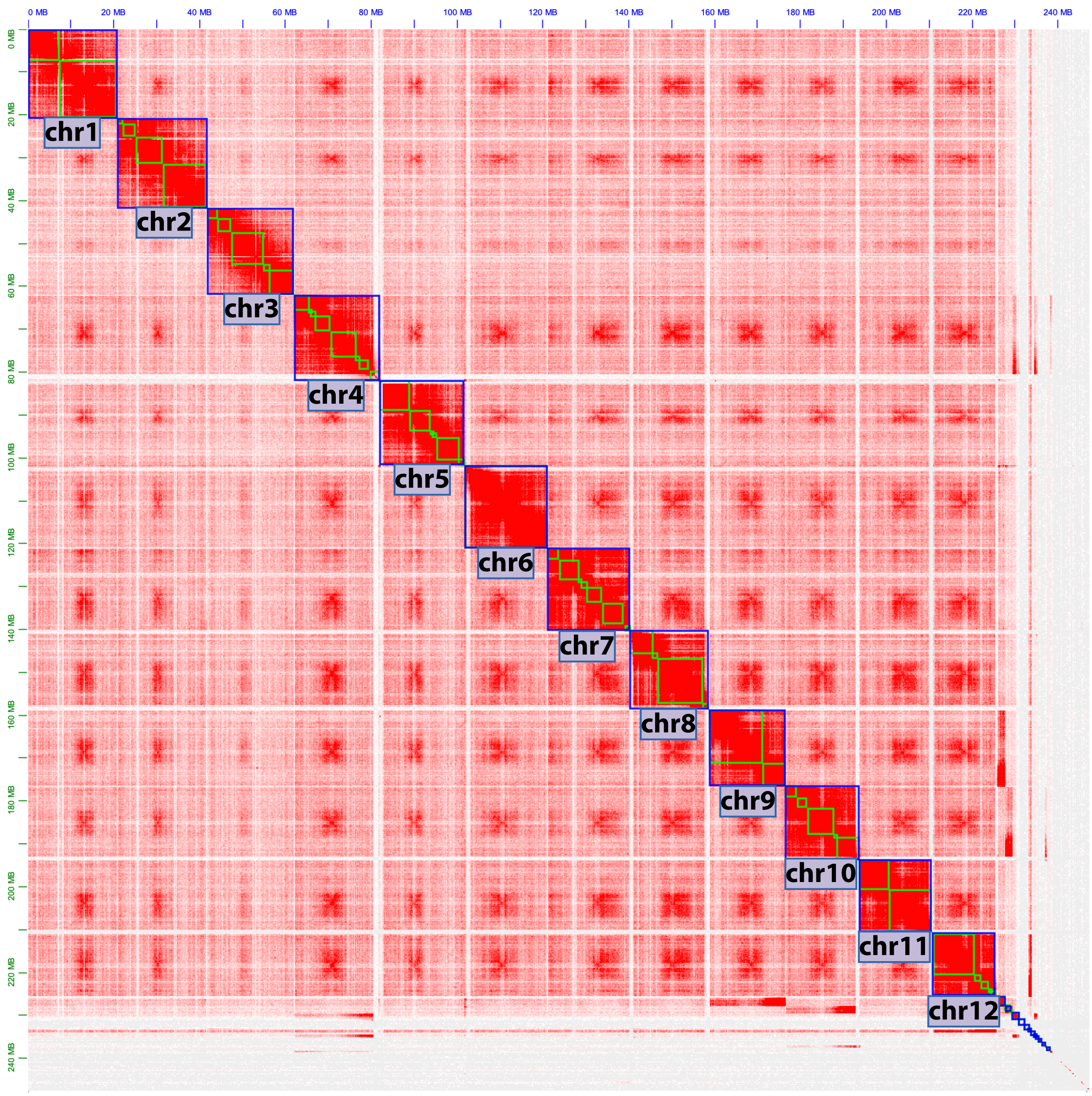

**Supplemental Figure 1.** Contact map illustrating the interaction frequency between HiC reads across the 12 major scaffolds of the *Tigriopus* aff. *angulatus* (qhTigAngs1.1.pri) assembly.

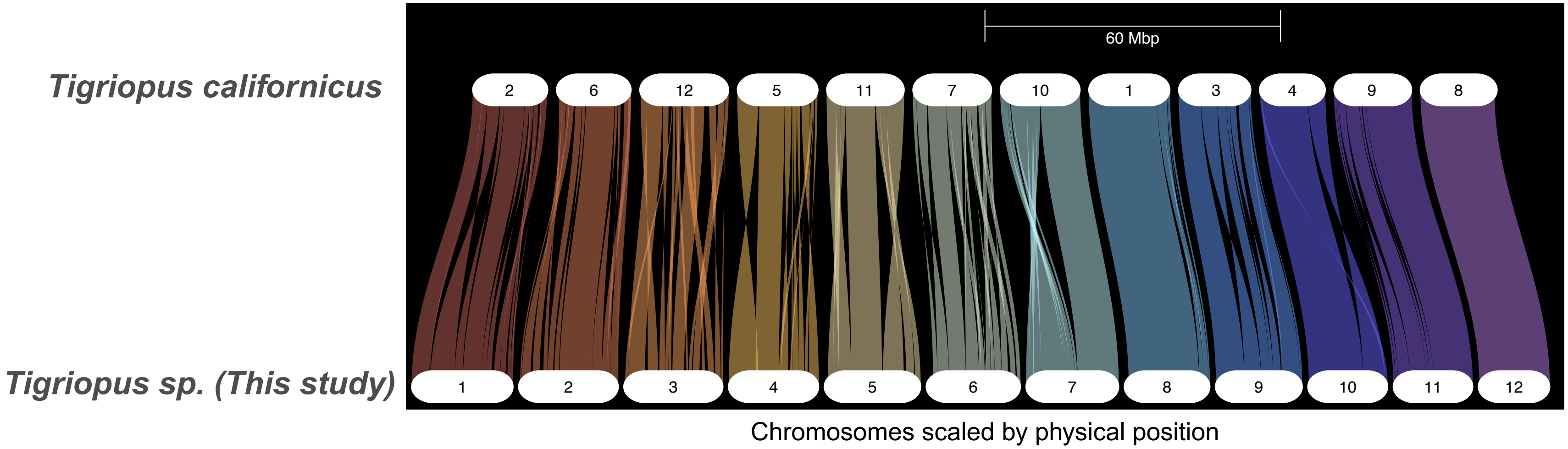

**Supplemental Figure 2.** Synteny in gene order between the assembly representing *Tigriopus* aff. *angulatus* (qhTigAngs1.1.pri) and *T. californicus* (GCF_007210705.1).

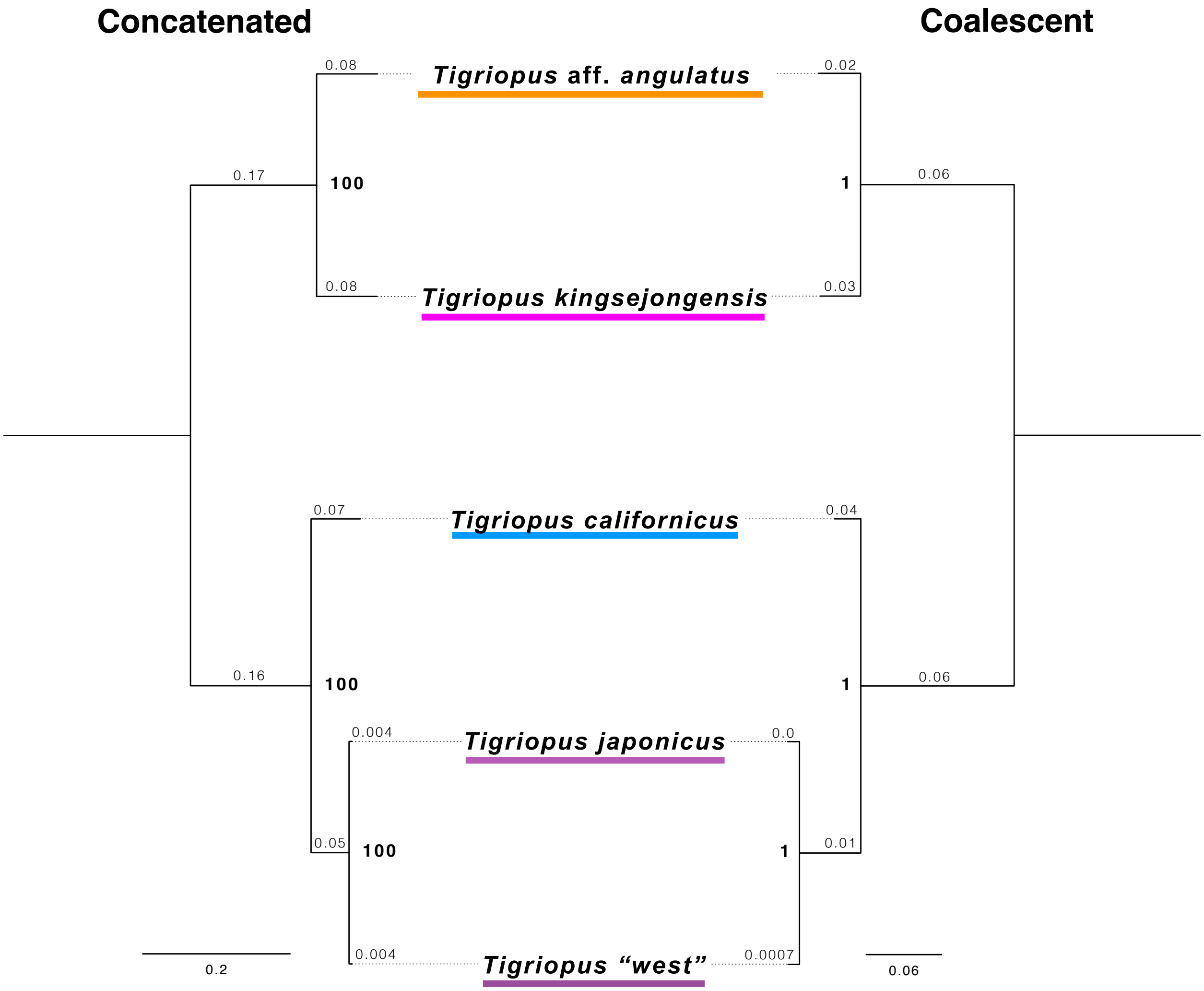

**Supplemental Figure 3.** Concatenated maximum likelihood and coalescent phylogenies inferred from 979 BUSCO genes extracted from the genome assemblies for each taxon. Bold numbers at the nodes indicate bootstrap (concatenated analysis) or local branch support, and numbers above the branches show branch length in number of substitutions per site. A Newick version of this tree is available in the supplemental files.

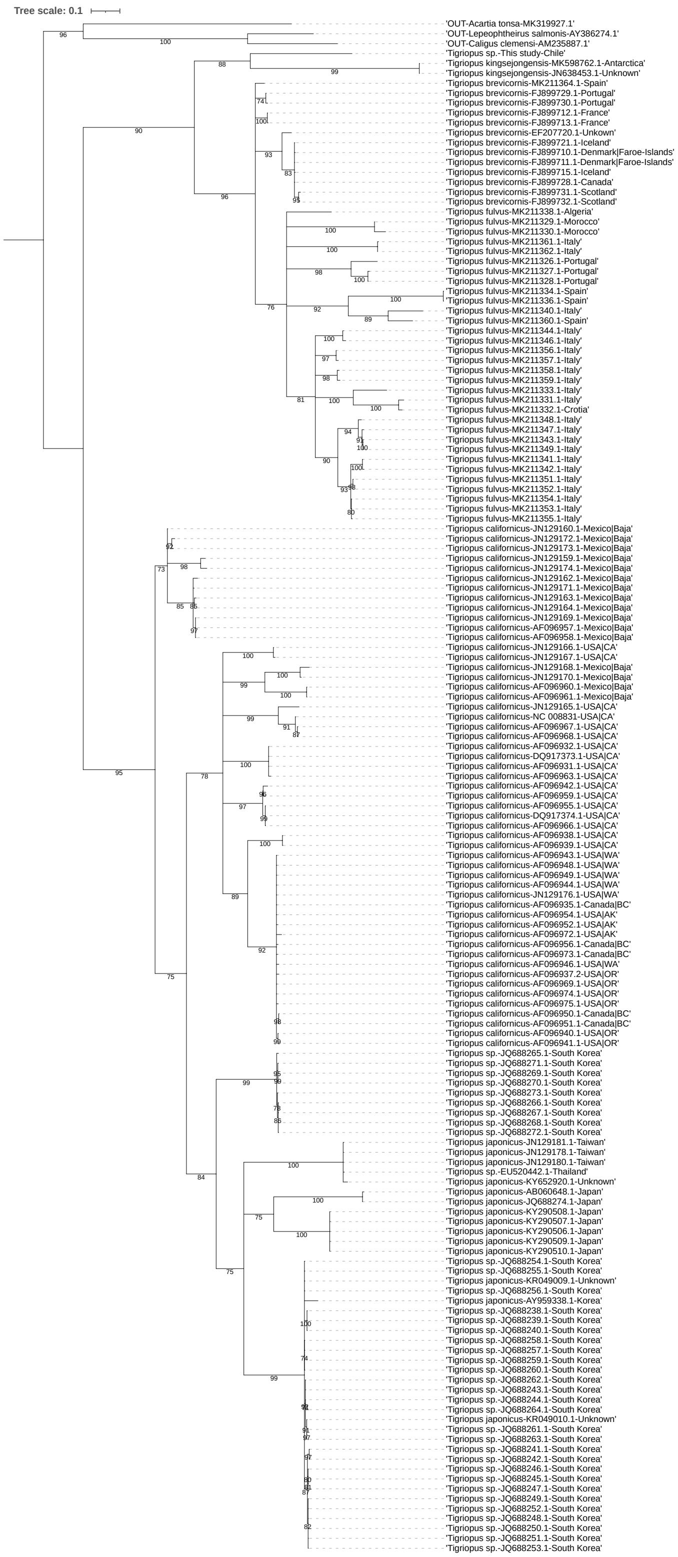

**Supplemental Figure 4**. Phylogeny of *Tigriopus* inferred from unique COXI sequences. Values at the nodes show bootstrap support, and bootstrap support values <70% have been collapsed. Additional details on sequences used are provided in Supplemental Table 7 and a Newick version of this tree is available in the supplemental files

**Supplemental Table 1.** Genome assembly statistics and BUSCO scores at different stages of the assembly process showing the effects of two rounds of contaminant removal (- cleanN) and two rounds of duplicate purging (- purgeN). The last column shows assembly statistics for the final, primary assembly after scaffolding.

| | Contigs | Contigs - clean1 | Contigs - clean2 | Contigs - purge1 | Contigs - purge2 | qhTigAngs1.1.pri |
|---|---|---|---|---|---|---|
| # scaffolds | 819 | 591 | 546 | 158 | 141 | 110 |
| Total scaffold length | 358,622,197 | 327,816,855 | 324,639,169 | 253,939,891 | 247,219,408 | 247,224,008 |
| Average scaffold length | 437,878 | 554,682 | 594,577 | 1,607,215 | 1,753,329 | 2,247,491 |
| Scaffold N50 | 4,863,442 | 5,265,128 | 5,265,128 | 6,322,232 | 6,924,626 | 19,081,170 |
| Scaffold auN | 6,093,784 | 6,461,798 | 6,522,864 | 8,065,791 | 8,215,725 | 17,378,083 |
| Scaffold L50 | 21 | 18 | 18 | 12 | 11 | 7 |
| Largest scaffold | 19,536,827 | 19,536,827 | 19,536,827 | 19,536,827 | 19,536,827 | 20,910,153 |
| Smallest scaffold | 7,734 | 9,044 | 9,044 | 14,431 | 14,431 | 14,431 |
| # contigs | 819 | 591 | 546 | 158 | 141 | 156 |
| Total contig length | 358,622,197 | 327,816,855 | 324,639,169 | 253,939,891 | 247,219,408 | 247,219,408 |
| Average contig length | 437,878 | 554,682 | 594,577 | 1,607,215 | 1,753,329 | 1,584,740 |
| Contig N50 | 4,863,442 | 5,265,128 | 5,265,128 | 6,322,232 | 6,924,626 | 6,305,818 |
| Contig auN | 6,093,784 | 6,461,798 | 6,522,864 | 8,065,791 | 8,215,725 | 7,280,570 |
| Contig L50 | 21 | 18 | 18 | 12 | 11 | 13 |
| Largest contig | 19,536,827 | 19,536,827 | 19,536,827 | 19,536,827 | 19,536,827 | 19,397,827 |
| Smallest contig | 7734 | 9044 | 9044 | 14431 | 14,431 | 14,431 |
| # gaps in scaffolds | 0 | 0 | 0 | 0 | 0 | 46 |
| Total gap length in scaffolds | 0 | 0 | 0 | 0 | 0 | 4,600 |
| Average gap length in scaffolds | 0 | 0 | 0 | 0 | 0 | 100 |
| Gap N50 in scaffolds | 0 | 0 | 0 | 0 | 0 | 100 |
| Gap auN in scaffolds | 0 | 0 | 0 | 0 | 0 | 100 |
| Gap L50 in scaffolds | 0 | 0 | 0 | 0 | 0 | 23 |
| Largest gap in scaffolds | 0 | 0 | 0 | 0 | 0 | 100 |
| Smallest gap in scaffolds | 0 | 0 | 0 | 0 | 0 | 100 |
| GC content % | 45.42 | 45.05 | 45.04 | 45.01 | 45.02 | 45 |
| # soft-masked bases | 0 | 0 | 0 | 0 | 0 | - |
| # segments | 819 | 591 | 546 | 158 | 141 | 156 |
| Total segment length | 358,622,197 | 327,816,855 | 324,639,169 | 253,939,891 | 247,219,408 | 247,219,408 |
| Average segment length | 437878.14 | 554,682 | 594,577 | 1,607,215 | 1,753,329 | 1,584,740 |
| # gaps | 0 | 0 | 0 | 0 | 0 | 46 |
| # paths | 819 | 591 | 546 | 158 | 141 | 110 |
| Single Copy Complete | | | S:72.06%, 730 | S:93.39%, 946 | S:96.64%, 979 | S:96.64%, 979 |
| Duplicated Complete | | | D:24.98%, 253 | D:3.75%, 38 | D:0.49%, 5 | D:0.49%, 5 |
| Fragmented | | | F:0.69%, 7 | F:0.69%, 7 | F:0.69%, 7 | F:0.69%, 7 |
| Missing | | | M:2.27%, 23 | M:2.17%, 22 | M:2.17%, 22 | M:2.17%, 22 |
| Count of Genes in Arthopod OrthoDBv10 | | | N:1013 | N:1013 | N:1013 | N:1013 |

**Supplemental Table 2.** Repeat classes and counts for the qhTigAngs1.1.pri assembly.

| | | | Number | Length (bp) | % of sequence |
|---|---|---|---|---|---|
| **Retroelements** | | | **84,911** | **68,970,554** | **27.9** |
| SINEs: | | | - | - | - |
| Penelope: | | | - | - | - |
| LINEs: | | | 39,153 | 35,226,231 | 14.3 |
| | CRE/SLACS | | - | - | - |
| | L2/CR1/Rex | | 33,014 | 25,792,917 | 10.4 |
| | R1/LOA/Jockey | | 106 | 76,377 | 0.03 |
| | R2/R4/NeSL | | - | - | - |
| | RTE/Bov-B | | - | - | - |
| | L1/CIN4 | | - | - | - |
| LTR elements: | | | 45,758 | 33,744,323 | 13.7 |
| | BEL/Pao | | 2,361 | 3,059,789 | 1.2 |
| | Ty1/Copia | | 1,106 | 486,716 | 0.2 |
| | Gypsy/DIRS1 | | 26,475 | 26,334,556 | 10.7 |
| | | Retroviral | - | - | - |
| **DNA transposons** | | | **13,106** | **5,182,429** | **2.1** |
| hobo-Activator | | | 1,078 | 340,137 | 0.1 |
| Tc1-IS630-Pogo | | | 8,994 | 3,603,548 | 1.5 |
| En-Spm | | | - | - | - |
| MULE-MuDR | | | 74 | 27,161 | 0.01 |
| PiggyBac | | | - | - | - |
| Tourist/Harbinger | | | - | - | - |
| Other (Mirage, P-element, Transib) | | | 1,795 | 627,022 | 0.3 |
| **Rolling-circles** | | | - | - | - |
| **Unclassified:** | | | **126,022** | **33,253,916** | **13.5** |
| **Total interspersed repeats:** | | | | **107,406,899** | **43.4** |
| **Small RNA:** | | | **1,016** | **278,242** | **0.1** |
| **Satellites:** | | | **1,049** | **346,211** | **0.1** |
| **Simple repeats:** | | | **14,071** | **1,161,081** | **0.5** |
| **Low complexity:** | | | **1,571** | **72,786** | **0.0** |

**Supplemental Table 3.** Summary statistics from AGAT (agat_sp_statistics.pl) for the protein coding gene models predicted by the NCBI egapx and EviAnn workflows using the qhTigAngs1.1.pri assembly.

| | egapx | EviAnn |
|---|---|---|
| Number of gene | 14,839 | 14,261 |
| Number of pseudogene | 160 | - |
| Number of mRNA | 22,383 | 29,129 |
| Number of mRNAs with utr both sides | 20,575 | 24,674 |
| Number of mRNAs with at least one utr | 21,734 | 27,448 |
| Number of cds | 22,188 | 29,074 |
| Number of exon | 142,222 | 179,191 |
| Number of five_prime_utr | 21,290 | 27,294 |
| Number of three_prime_utr | 21,019 | 24,828 |
| Number of exon in cds | 131,672 | 159,029 |
| Number of exon in five_prime_utr | 30,345 | 42,359 |
| Number of exon in three_prime_utr | 22,017 | 29,692 |
| Number of intron in cds | 109,484 | 129,955 |
| Number of intron in exon | 119,839 | 150,062 |
| Number of intron in five_prime_utr | 9,055 | 15,065 |
| Number of intron in three_prime_utr | 998 | 4,864 |
| Number gene..pseudogene overlapping | 1,476 | - |
| Number gene overlapping | - | 1,597 |
| Number of single exon gene | 1,323 | 2,177 |
| Number of single exon pseudogene | 100 | - |
| Number of single exon mRNA | 1,427 | 2,893 |
| Mean mRNAs per gene | 2 | 2 |
| Mean mRNAs per pseudogene | 140 | - |
| Mean cdss per mRNA | 1 | 1 |
| Mean exons per mRNA | 6 | 6 |
| Mean five_prime_utrs per mRNA | 1 | 1 |
| Mean three_prime_utrs per mRNA | 1 | 1 |
| Mean exons per cds | 6 | 6 |
| Mean exons per five_prime_utr | 1 | 2 |
| Mean exons per three_prime_utr | 1 | 1 |
| Mean introns in cdss per mRNA | 5 | 5 |
| Mean introns in exons per mRNA | 5 | 5 |
| Mean introns in five_prime_utrs per mRNA | 0 | 1 |
| Mean introns in three_prime_utrs per mRNA | - | 0 |
| Total gene length (bp) | 128,496,774 | 122,264,036 |
| Total pseudogene length (bp) | 383,319 | - |
| Total mRNA length (bp) | 255,428,913 | 308,625,198 |
| Total cds length (bp) | 42,469,014 | 51,956,640 |
| Total exon length (bp) | 56,880,451 | 78,190,606 |
| Total five_prime_utr length (bp) | 7,430,061 | 12,639,346 |
| Total three_prime_utr length (bp) | 6,608,820 | 13,529,780 |
| Total intron length per cds (bp) | 159,437,714 | 179,670,901 |
| Total intron length per exon (bp) | 198,548,462 | 230,434,592 |
| Total intron length per five_prime_utr (bp) | 36,896,098 | 45,578,062 |
| Total intron length per three_prime_utr (bp) | 1,341,947 | 4,682,515 |
| Mean gene length (bp) | 8,659 | 8,573 |
| Mean pseudogene length (bp) | 2,396 | - |
| Mean mRNA length (bp) | 11,412 | 10,595 |
| Mean cds length (bp) | 1,914 | 1,787 |
| Mean exon length (bp) | 400 | 436 |
| Complete BUSCOs (C) | 96.7%, 979 | 95.4%, 966 |
| Complete and single-copy BUSCOs (S) | 95.1%, 963 | 93.8%, 950 |
| Complete and duplicated BUSCOs (D) | 1.6%, 16 | 1.6%, 16 |
| Fragmented BUSCOs (F) | 0.2%, 2 | 0.7%, 7 |
| Missing BUSCOs (M) | 3.1%, 32 | 3.9%, 40 |
| Total BUSCOs | 1,013.00 | 1,013 |

**Supplemental Table 4.** Comparison of assembly and completeness metrics for the qhTigAngs1.1.pri assembly compared to genome assemblies for other *Tigriopus* taxa that are publicly available as of June 2026.

| | qhTigAngs1.1.pri (This study) | *T. californicus - GCF_007210705.1* | *T. japonicus - GCA_010645155.1* | *T. kingsejongensis - GCA_012959195.1* | *T. "West" - GCA_051201535.1* |
|---|---|---|---|---|---|
| # scaffolds | 111 | 459 | 339 | 938 | 1,997 |
| Total scaffold length | 247,240,542 | 191,142,546 | 196,599,007 | 338,647,387 | 176,799,855 |
| Average scaffold length | 2,227,392 | 416,433 | 579,938 | 361,031 | 88,533 |
| Scaffold N50 | 19,081,170 | 15,806,032 | 10,654,335 | 1,473,880 | 11,193,223 |
| Scaffold auN | 17,376,922 | 15,777,613 | 8,683,405 | 2,224,281 | 9,169,004 |
| Scaffold L50 | 7 | 6 | 8 | 55 | 7 |
| Largest scaffold | 20,910,153 | 18,073,795 | 15,288,102 | 9,103,457 | 13,520,769 |
| Smallest scaffold | 14,431 | 734 | 1,000 | 1,000 | 10,000 |
| # contigs | 157 | 14,465 | 6,805 | 1,097 | 6,617 |
| Total contig length | 247,235,942 | 187,155,163 | 191,613,931 | 338,102,786 | 176,337,855 |
| Average contig length | 1,574,751 | 12,938 | 28,158 | 308,207 | 26,649 |
| Contig N50 | 6,305,818 | 34,267 | 69,577 | 1,293,995 | 36,733 |
| Contig auN | 7,280,084 | 42,646 | 96,141 | 2,099,008 | 47,088 |
| Contig L50 | 13 | 1,621 | 749 | 62 | 1,439 |
| Largest contig | 19,397,827 | 210,266 | 529,501 | 9,103,457 | 279,458 |
| Smallest contig | 14,431 | 1 | 5 | 2 | 1,009 |
| # gaps in scaffolds | 46 | 14,006 | 6,466 | 159 | 4,620 |
| Total gap length in scaffolds | 4,600 | 3,987,383 | 4,985,076 | 544,601 | 462,000 |
| Average gap length in scaffolds | 100 | 285 | 771 | 3,425 | 100 |
| Gap N50 in scaffolds | 100 | 1,718 | 2,675 | 4,726 | 100 |
| Gap auN in scaffolds | 100 | 1,836 | 3,236 | 6,387 | 100 |
| Gap L50 in scaffolds | 23 | 759 | 532 | 32 | 2,310 |
| Largest gap in scaffolds | 100 | 5,651 | 9,044 | 22,889 | 100 |
| Smallest gap in scaffolds | 100 | 1 | 1 | 51 | 100 |
| GC content % | 45 | 42 | 40 | 47 | 40 |
| # soft-masked bases | 109,265,219 | 33,788,852 | 45,144,196 | 55,464,373 | 36,952,373 |
| # segments | 157 | 14,465 | 6,805 | 1,097 | 6,617 |
| Total segment length | 247,235,942 | 187,155,162 | 191,613,931 | 338,102,786 | 176,337,855 |
| Average segment length | 1,574,751 | 12,938 | 28,158 | 308,207 | 26,649 |
| # gaps | 46 | 14,006 | 6,466 | 159 | 4,620 |
| # paths | 111 | 459 | 339 | 938 | 1,997 |
| Single Copy Complete | S:95.2%, 964 | S:95.6%, 968 | S:95.1%, 963 | S:94.9%, 961 | S:86.9%, 880 |
| Duplicated Complete | D:1.7%, 17 | D:1.0%, 10 | D:1.4%, 14 | D:2.2%, 22 | D:5.5%, 56 |
| Fragmented | F:0.7%, 7 | F:1.2%, 12 | F:1.4%, 14 | F:0.8%, 8 | F:2.1%, 21 |
| Missing | M:2.4%, 25 | M:2.2%, 23 | M:2.1%, 22 | M:2.1%, 22 | M:5.5%, 56 |
| Count of Genes in Arthopod OrthoDBv10 | N:1013 | N:1013 | N:1013 | N:1013 | N:1013 |

**Supplementary Table 5.** Evolutionary distances between COX1 sequences of *Tigriopus* aff. *angulatus* in this study and other *Tigriopus* species computed as the proportion of nucleotide sites at which they were different (p-distance) using MEGA 12.1. Taxon names include the NCBI accession number of the sequence used and the country in which the sample was collected.

| | *T. californicus* - NC_008831.2 - USA\|CA | *T. japonicus* - AB060648.1 - Japan | *T. kingsejongensis* - MK598762.1 - Antarctica |
|---|---|---|---|
| *T. japonicus* - AB060648.1 - Japan | 0.23 | - | - |
| *T. kingsejongensis* - MK598762.1 - Antarctica | 0.32 | 0.33 | - |
| *Tigriopus* aff. *angulatus* - This study - Chile | 0.31 | 0.30 | 0.26 |

**Supplementary Table 6.** Evolutionary distances between COX1 sequences of *Tigriopus* aff. *angulatus* in this study and other Tigriopus species computed using the Jukes-Cantor model (Jukes and Cantor 1969) in MEGA 12.1. Taxon names include the NCBI accession number of the sequence used and the country in which the sample was collected.

| | *T. californicus* - NC_008831.2 - USA\|CA | *T. japonicus* - AB060648.1 - Japan | *T. kingsejongensis* - MK598762.1 - Antarctica |
|---|---|---|---|
| *T. japonicus* - AB060648.1 - Japan | 0.27 | - | - |
| *T. kingsejongensis* - MK598762.1 - Antarctica | 0.41 | 0.43 | - |
| *Tigriopus* aff. *angulatus* - This study - Chile | 0.40 | 0.38 | 0.32 |

**Supplementary Table 7.** Unique COX1 sequences downloaded from NCBI GenBank and used to infer a mitochondrial phylogeny of *Tigriopus* (Figure 2, Supplemental Figure 4).

| Taxon | NCBI GenBank Accession | Country of Collection | Tip Name in Phylogeny | Reference (DOI) |
|---|---|---|---|---|
| Tigriopus brevicornis | EF207720.1 | Unkown | Tigriopus brevicornis-EF207720.1-Unkown | 10.3989/scimar.2009.73n3579 |
| Tigriopus brevicornis | FJ899710.1 | Denmark (Faroe Islands) | Tigriopus brevicornis-FJ899710.1-Denmark (Faroe Islands) | 10.1007/s00227-010-1415-7 |
| Tigriopus brevicornis | FJ899711.1 | Denmark (Faroe Islands) | Tigriopus brevicornis-FJ899711.1-Denmark (Faroe Islands) | 10.1007/s00227-010-1415-7 |
| Tigriopus brevicornis | FJ899712.1 | France | Tigriopus brevicornis-FJ899712.1-France | 10.1007/s00227-010-1415-7 |
| Tigriopus brevicornis | FJ899713.1 | France | Tigriopus brevicornis-FJ899713.1-France | 10.1007/s00227-010-1415-7 |
| Tigriopus brevicornis | FJ899715.1 | Iceland | Tigriopus brevicornis-FJ899715.1-Iceland | 10.1007/s00227-010-1415-7 |
| Tigriopus brevicornis | FJ899721.1 | Iceland | Tigriopus brevicornis-FJ899721.1-Iceland | 10.1007/s00227-010-1415-7 |
| Tigriopus brevicornis | FJ899728.1 | Canada | Tigriopus brevicornis-FJ899728.1-Canada | 10.1007/s00227-010-1415-7 |
| Tigriopus brevicornis | FJ899729.1 | Portugal | Tigriopus brevicornis-FJ899729.1-Portugal | 10.1007/s00227-010-1415-7 |
| Tigriopus brevicornis | FJ899730.1 | Portugal | Tigriopus brevicornis-FJ899730.1-Portugal | 10.1007/s00227-010-1415-7 |
| Tigriopus brevicornis | FJ899731.1 | Scotland | Tigriopus brevicornis-FJ899731.1-Scotland | 10.1007/s00227-010-1415-7 |
| Tigriopus brevicornis | FJ899732.1 | Scotland | Tigriopus brevicornis-FJ899732.1-Scotland | 10.1007/s00227-010-1415-7 |
| Tigriopus brevicornis | MK211364.1 | Spain | Tigriopus brevicornis-MK211364.1-Spain | 10.7773/cm.v45i2.2946 |
| Tigriopus californicus | AF096931.1 | USA:CA | Tigriopus californicus-AF096931.1-USA:CA | 10.1046/j.0962-1083.2001.01306.x |
| Tigriopus californicus | AF096932.1 | USA:CA | Tigriopus californicus-AF096932.1-USA:CA | 10.1046/j.0962-1083.2001.01306.x |
| Tigriopus californicus | AF096935.1 | Canada:BC | Tigriopus californicus-AF096935.1-Canada:BC | 10.1046/j.0962-1083.2001.01306.x |
| Tigriopus californicus | AF096937.2 | USA:OR | Tigriopus californicus-AF096937.2-USA:OR | 10.1046/j.0962-1083.2001.01306.x |
| Tigriopus californicus | AF096938.1 | USA:CA | Tigriopus californicus-AF096938.1-USA:CA | 10.1046/j.0962-1083.2001.01306.x |
| Tigriopus californicus | AF096939.1 | USA:CA | Tigriopus californicus-AF096939.1-USA:CA | 10.1046/j.0962-1083.2001.01306.x |
| Tigriopus californicus | AF096940.1 | USA:OR | Tigriopus californicus-AF096940.1-USA:OR | 10.1046/j.0962-1083.2001.01306.x |
| Tigriopus californicus | AF096941.1 | USA:OR | Tigriopus californicus-AF096941.1-USA:OR | 10.1046/j.0962-1083.2001.01306.x |
| Tigriopus californicus | AF096942.1 | USA:CA | Tigriopus californicus-AF096942.1-USA:CA | 10.1046/j.0962-1083.2001.01306.x |
| Tigriopus californicus | AF096943.1 | USA:WA | Tigriopus californicus-AF096943.1-USA:WA | 10.1046/j.0962-1083.2001.01306.x |
| Tigriopus californicus | AF096944.1 | USA:WA | Tigriopus californicus-AF096944.1-USA:WA | 10.1046/j.0962-1083.2001.01306.x |
| Tigriopus californicus | AF096946.1 | USA:WA | Tigriopus californicus-AF096946.1-USA:WA | 10.1046/j.0962-1083.2001.01306.x |
| Tigriopus californicus | AF096948.1 | USA:WA | Tigriopus californicus-AF096948.1-USA:WA | 10.1046/j.0962-1083.2001.01306.x |
| Tigriopus californicus | AF096949.1 | USA:WA | Tigriopus californicus-AF096949.1-USA:WA | 10.1046/j.0962-1083.2001.01306.x |
| Tigriopus californicus | AF096950.1 | Canada:BC | Tigriopus californicus-AF096950.1-Canada:BC | 10.1046/j.0962-1083.2001.01306.x |
| Tigriopus californicus | AF096951.1 | Canada:BC | Tigriopus californicus-AF096951.1-Canada:BC | 10.1046/j.0962-1083.2001.01306.x |
| Tigriopus californicus | AF096952.1 | USA:AK | Tigriopus californicus-AF096952.1-USA:AK | 10.1046/j.0962-1083.2001.01306.x |
| Tigriopus californicus | AF096954.1 | USA:AK | Tigriopus californicus-AF096954.1-USA:AK | 10.1046/j.0962-1083.2001.01306.x |
| Tigriopus californicus | AF096955.1 | USA:CA | Tigriopus californicus-AF096955.1-USA:CA | 10.1046/j.0962-1083.2001.01306.x |

| | | | | |
|---|---|---|---|---|
| Tigriopus californicus | AF096956.1 | Canada:BC | Tigriopus californicus-AF096956.1-Canada:BC | 10.1046/j.0962-1083.2001.01306.x |
| Tigriopus californicus | AF096957.1 | Mexico:Baja | Tigriopus californicus-AF096957.1-Mexico:Baja | 10.1046/j.0962-1083.2001.01306.x |
| Tigriopus californicus | AF096958.1 | Mexico:Baja | Tigriopus californicus-AF096958.1-Mexico:Baja | 10.1046/j.0962-1083.2001.01306.x |
| Tigriopus californicus | AF096959.1 | USA:CA | Tigriopus californicus-AF096959.1-USA:CA | 10.1046/j.0962-1083.2001.01306.x |
| Tigriopus californicus | AF096960.1 | Mexico:Baja | Tigriopus californicus-AF096960.1-Mexico:Baja | 10.1046/j.0962-1083.2001.01306.x |
| Tigriopus californicus | AF096961.1 | Mexico:Baja | Tigriopus californicus-AF096961.1-Mexico:Baja | 10.1046/j.0962-1083.2001.01306.x |
| Tigriopus californicus | AF096963.1 | USA:CA | Tigriopus californicus-AF096963.1-USA:CA | 10.1046/j.0962-1083.2001.01306.x |
| Tigriopus californicus | AF096966.1 | USA:CA | Tigriopus californicus-AF096966.1-USA:CA | 10.1046/j.0962-1083.2001.01306.x |
| Tigriopus californicus | AF096967.1 | USA:CA | Tigriopus californicus-AF096967.1-USA:CA | 10.1046/j.0962-1083.2001.01306.x |
| Tigriopus californicus | AF096968.1 | USA:CA | Tigriopus californicus-AF096968.1-USA:CA | 10.1046/j.0962-1083.2001.01306.x |
| Tigriopus californicus | AF096969.1 | USA:OR | Tigriopus californicus-AF096969.1-USA:OR | 10.1046/j.0962-1083.2001.01306.x |
| Tigriopus californicus | AF096972.1 | USA:AK | Tigriopus californicus-AF096972.1-USA:AK | 10.1046/j.0962-1083.2001.01306.x |
| Tigriopus californicus | AF096973.1 | Canada:BC | Tigriopus californicus-AF096973.1-Canada:BC | 10.1046/j.0962-1083.2001.01306.x |
| Tigriopus californicus | AF096974.1 | USA:OR | Tigriopus californicus-AF096974.1-USA:OR | 10.1046/j.0962-1083.2001.01306.x |
| Tigriopus californicus | AF096975.1 | USA:OR | Tigriopus californicus-AF096975.1-USA:OR | 10.1046/j.0962-1083.2001.01306.x |
| Tigriopus californicus | DQ917373.1 | USA:CA | Tigriopus californicus-DQ917373.1-USA:CA | 10.1016/j.gene.2007.07.026 |
| Tigriopus californicus | DQ917374.1 | USA:CA | Tigriopus californicus-DQ917374.1-USA:CA | 10.1016/j.gene.2007.07.026 |
| Tigriopus californicus | JN129159.1 | Mexico:Baja | Tigriopus californicus-JN129159.1-Mexico:Baja | 10.1111/jbi.12107 |
| Tigriopus californicus | JN129160.1 | Mexico:Baja | Tigriopus californicus-JN129160.1-Mexico:Baja | 10.1111/jbi.12107 |
| Tigriopus californicus | JN129162.1 | Mexico:Baja | Tigriopus californicus-JN129162.1-Mexico:Baja | 10.1111/jbi.12107 |
| Tigriopus californicus | JN129163.1 | Mexico:Baja | Tigriopus californicus-JN129163.1-Mexico:Baja | 10.1111/jbi.12107 |
| Tigriopus californicus | JN129164.1 | Mexico:Baja | Tigriopus californicus-JN129164.1-Mexico:Baja | 10.1111/jbi.12107 |
| Tigriopus californicus | JN129165.1 | USA:CA | Tigriopus californicus-JN129165.1-USA:CA | 10.1111/jbi.12107 |
| Tigriopus californicus | JN129166.1 | USA:CA | Tigriopus californicus-JN129166.1-USA:CA | 10.1111/jbi.12107 |
| Tigriopus californicus | JN129167.1 | USA:CA | Tigriopus californicus-JN129167.1-USA:CA | 10.1111/jbi.12107 |
| Tigriopus californicus | JN129168.1 | Mexico:Baja | Tigriopus californicus-JN129168.1-Mexico:Baja | 10.1111/jbi.12107 |
| Tigriopus californicus | JN129169.1 | Mexico:Baja | Tigriopus californicus-JN129169.1-Mexico:Baja | 10.1111/jbi.12107 |
| Tigriopus californicus | JN129170.1 | Mexico:Baja | Tigriopus californicus-JN129170.1-Mexico:Baja | 10.1111/jbi.12107 |
| Tigriopus californicus | JN129171.1 | Mexico:Baja | Tigriopus californicus-JN129171.1-Mexico:Baja | 10.1111/jbi.12107 |
| Tigriopus californicus | JN129172.1 | Mexico:Baja | Tigriopus californicus-JN129172.1-Mexico:Baja | 10.1111/jbi.12107 |
| Tigriopus californicus | JN129173.1 | Mexico:Baja | Tigriopus californicus-JN129173.1-Mexico:Baja | 10.1111/jbi.12107 |
| Tigriopus californicus | JN129174.1 | Mexico:Baja | Tigriopus californicus-JN129174.1-Mexico:Baja | 10.1111/jbi.12107 |
| Tigriopus californicus | JN129176.1 | USA:WA | Tigriopus californicus-JN129176.1-USA:WA | 10.1111/jbi.12107 |
| Tigriopus californicus | NC_008831.2 | USA:CA | Tigriopus californicus-NC_008831-USA:CA | 10.1016/j.gene.2007.07.026 |
| Tigriopus fulvus | MK211326.1 | Portugal | Tigriopus fulvus-MK211326.1-Portugal | 10.7773/cm.v45i2.2946 |
| Tigriopus fulvus | MK211327.1 | Portugal | Tigriopus fulvus-MK211327.1-Portugal | 10.7773/cm.v45i2.2946 |

| | | | | |
|---|---|---|---|---|
| Tigriopus fulvus | MK211328.1 | Portugal | Tigriopus fulvus-MK211328.1-Portugal | 10.7773/cm.v45i2.2946 |
| Tigriopus fulvus | MK211329.1 | Morocco | Tigriopus fulvus-MK211329.1-Morocco | 10.7773/cm.v45i2.2946 |
| Tigriopus fulvus | MK211330.1 | Morocco | Tigriopus fulvus-MK211330.1-Morocco | 10.7773/cm.v45i2.2946 |
| Tigriopus fulvus | MK211331.1 | Italy | Tigriopus fulvus-MK211331.1-Italy | 10.7773/cm.v45i2.2946 |
| Tigriopus fulvus | MK211332.1 | Crotia | Tigriopus fulvus-MK211332.1-Crotia | 10.7773/cm.v45i2.2946 |
| Tigriopus fulvus | MK211333.1 | Italy | Tigriopus fulvus-MK211333.1-Italy | 10.7773/cm.v45i2.2946 |
| Tigriopus fulvus | MK211334.1 | Spain | Tigriopus fulvus-MK211334.1-Spain | 10.7773/cm.v45i2.2946 |
| Tigriopus fulvus | MK211336.1 | Spain | Tigriopus fulvus-MK211336.1-Spain | 10.7773/cm.v45i2.2946 |
| Tigriopus fulvus | MK211338.1 | Algeria | Tigriopus fulvus-MK211338.1-Algeria | 10.7773/cm.v45i2.2946 |
| Tigriopus fulvus | MK211340.1 | Italy | Tigriopus fulvus-MK211340.1-Italy | 10.7773/cm.v45i2.2946 |
| Tigriopus fulvus | MK211341.1 | Italy | Tigriopus fulvus-MK211341.1-Italy | 10.7773/cm.v45i2.2946 |
| Tigriopus fulvus | MK211342.1 | Italy | Tigriopus fulvus-MK211342.1-Italy | 10.7773/cm.v45i2.2946 |
| Tigriopus fulvus | MK211343.1 | Italy | Tigriopus fulvus-MK211343.1-Italy | 10.7773/cm.v45i2.2946 |
| Tigriopus fulvus | MK211344.1 | Italy | Tigriopus fulvus-MK211344.1-Italy | 10.7773/cm.v45i2.2946 |
| Tigriopus fulvus | MK211346.1 | Italy | Tigriopus fulvus-MK211346.1-Italy | 10.7773/cm.v45i2.2946 |
| Tigriopus fulvus | MK211347.1 | Italy | Tigriopus fulvus-MK211347.1-Italy | 10.7773/cm.v45i2.2946 |
| Tigriopus fulvus | MK211348.1 | Italy | Tigriopus fulvus-MK211348.1-Italy | 10.7773/cm.v45i2.2946 |
| Tigriopus fulvus | MK211349.1 | Italy | Tigriopus fulvus-MK211349.1-Italy | 10.7773/cm.v45i2.2946 |
| Tigriopus fulvus | MK211351.1 | Italy | Tigriopus fulvus-MK211351.1-Italy | 10.7773/cm.v45i2.2946 |
| Tigriopus fulvus | MK211352.1 | Italy | Tigriopus fulvus-MK211352.1-Italy | 10.7773/cm.v45i2.2946 |
| Tigriopus fulvus | MK211353.1 | Italy | Tigriopus fulvus-MK211353.1-Italy | 10.7773/cm.v45i2.2946 |
| Tigriopus fulvus | MK211354.1 | Italy | Tigriopus fulvus-MK211354.1-Italy | 10.7773/cm.v45i2.2946 |
| Tigriopus fulvus | MK211355.1 | Italy | Tigriopus fulvus-MK211355.1-Italy | 10.7773/cm.v45i2.2946 |
| Tigriopus fulvus | MK211356.1 | Italy | Tigriopus fulvus-MK211356.1-Italy | 10.7773/cm.v45i2.2946 |
| Tigriopus fulvus | MK211357.1 | Italy | Tigriopus fulvus-MK211357.1-Italy | 10.7773/cm.v45i2.2946 |
| Tigriopus fulvus | MK211358.1 | Italy | Tigriopus fulvus-MK211358.1-Italy | 10.7773/cm.v45i2.2946 |
| Tigriopus fulvus | MK211359.1 | Italy | Tigriopus fulvus-MK211359.1-Italy | 10.7773/cm.v45i2.2946 |
| Tigriopus fulvus | MK211360.1 | Spain | Tigriopus fulvus-MK211360.1-Spain | 10.7773/cm.v45i2.2946 |
| Tigriopus fulvus | MK211361.1 | Italy | Tigriopus fulvus-MK211361.1-Italy | 10.7773/cm.v45i2.2946 |
| Tigriopus fulvus | MK211362.1 | Italy | Tigriopus fulvus-MK211362.1-Italy | 10.7773/cm.v45i2.2946 |
| Tigriopus japonicus | AB060648.1 | Japan | Tigriopus japonicus-AB060648.1-Japan | 10.1007/s10126-002-0033-x |
| Tigriopus japonicus | AY959338.1 | Korea | Tigriopus japonicus-AY959338.1-Korea | 10.1016/j.jembe.2005.12.047 |
| Tigriopus japonicus | JN129178.1 | Taiwan | Tigriopus japonicus-JN129178.1-Taiwan | 10.1111/jbi.12107 |
| Tigriopus japonicus | JN129180.1 | Taiwan | Tigriopus japonicus-JN129180.1-Taiwan | 10.1111/jbi.12107 |
| Tigriopus japonicus | JN129181.1 | Taiwan | Tigriopus japonicus-JN129181.1-Taiwan | 10.1111/jbi.12107 |
| Tigriopus japonicus | JQ688274.1 | Japan | Tigriopus japonicus-JQ688274.1-Japan | Unpublished |

| | | | | |
|---|---|---|---|---|
| Tigriopus japonicus | KR049009.1 | Unknown | Tigriopus japonicus-KR049009.1-Unknown | 10.1371/journal.pone.0157307 |
| Tigriopus japonicus | KR049010.1 | Unknown | Tigriopus japonicus-KR049010.1-Unknown | 10.1371/journal.pone.0157307 |
| Tigriopus japonicus | KY290506.1 | Japan | Tigriopus japonicus-KY290506.1-Japan | 10.1071/IS17002 |
| Tigriopus japonicus | KY290507.1 | Japan | Tigriopus japonicus-KY290507.1-Japan | 10.1071/IS17002 |
| Tigriopus japonicus | KY290508.1 | Japan | Tigriopus japonicus-KY290508.1-Japan | 10.1071/IS17002 |
| Tigriopus japonicus | KY290509.1 | Japan | Tigriopus japonicus-KY290509.1-Japan | 10.1071/IS17002 |
| Tigriopus japonicus | KY290510.1 | Japan | Tigriopus japonicus-KY290510.1-Japan | 10.1071/IS17002 |
| Tigriopus japonicus | KY652920.1 | Unknown | Tigriopus japonicus-KY652920.1-Unknown | Unpublished |
| Tigriopus kingsejongensis | JN638453.1 | Unknown | Tigriopus kingsejongensis-JN638453.1-Unknown | 10.2988/0006-324X-127.1.138 |
| Tigriopus kingsejongensis | MK598762.1 | Antarctica | Tigriopus kingsejongensis-MK598762.1-Antarctica | 10.1080/23802359.2019.1601042 |
| Tigriopus aff. angulatus | (This study) | Chile | Tigriopus sp.-This study-Chile | - |
| Tigriopus sp. | EU520442.1 | Thailand | Tigriopus sp.-EU520442.1-Thailand | Unpublished |
| Tigriopus sp. | JQ688238.1 | South Korea | Tigriopus sp.-JQ688238.1-South Korea | 10.1071/IS17002 |
| Tigriopus sp. | JQ688239.1 | South Korea | Tigriopus sp.-JQ688239.1-South Korea | 10.1071/IS17002 |
| Tigriopus sp. | JQ688240.1 | South Korea | Tigriopus sp.-JQ688240.1-South Korea | 10.1071/IS17002 |
| Tigriopus sp. | JQ688241.1 | South Korea | Tigriopus sp.-JQ688241.1-South Korea | 10.1071/IS17002 |
| Tigriopus sp. | JQ688242.1 | South Korea | Tigriopus sp.-JQ688242.1-South Korea | 10.1071/IS17002 |
| Tigriopus sp. | JQ688243.1 | South Korea | Tigriopus sp.-JQ688243.1-South Korea | 10.1071/IS17002 |
| Tigriopus sp. | JQ688244.1 | South Korea | Tigriopus sp.-JQ688244.1-South Korea | 10.1071/IS17002 |
| Tigriopus sp. | JQ688245.1 | South Korea | Tigriopus sp.-JQ688245.1-South Korea | 10.1071/IS17002 |
| Tigriopus sp. | JQ688246.1 | South Korea | Tigriopus sp.-JQ688246.1-South Korea | 10.1071/IS17002 |
| Tigriopus sp. | JQ688247.1 | South Korea | Tigriopus sp.-JQ688247.1-South Korea | 10.1071/IS17002 |
| Tigriopus sp. | JQ688248.1 | South Korea | Tigriopus sp.-JQ688248.1-South Korea | 10.1071/IS17002 |
| Tigriopus sp. | JQ688249.1 | South Korea | Tigriopus sp.-JQ688249.1-South Korea | 10.1071/IS17002 |
| Tigriopus sp. | JQ688250.1 | South Korea | Tigriopus sp.-JQ688250.1-South Korea | 10.1071/IS17002 |
| Tigriopus sp. | JQ688251.1 | South Korea | Tigriopus sp.-JQ688251.1-South Korea | 10.1071/IS17002 |
| Tigriopus sp. | JQ688252.1 | South Korea | Tigriopus sp.-JQ688252.1-South Korea | 10.1071/IS17002 |
| Tigriopus sp. | JQ688253.1 | South Korea | Tigriopus sp.-JQ688253.1-South Korea | 10.1071/IS17002 |
| Tigriopus sp. | JQ688254.1 | South Korea | Tigriopus sp.-JQ688254.1-South Korea | 10.1071/IS17002 |
| Tigriopus sp. | JQ688255.1 | South Korea | Tigriopus sp.-JQ688255.1-South Korea | 10.1071/IS17002 |
| Tigriopus sp. | JQ688256.1 | South Korea | Tigriopus sp.-JQ688256.1-South Korea | 10.1071/IS17002 |
| Tigriopus sp. | JQ688257.1 | South Korea | Tigriopus sp.-JQ688257.1-South Korea | 10.1071/IS17002 |
| Tigriopus sp. | JQ688258.1 | South Korea | Tigriopus sp.-JQ688258.1-South Korea | 10.1071/IS17002 |
| Tigriopus sp. | JQ688259.1 | South Korea | Tigriopus sp.-JQ688259.1-South Korea | 10.1071/IS17002 |
| Tigriopus sp. | JQ688260.1 | South Korea | Tigriopus sp.-JQ688260.1-South Korea | 10.1071/IS17002 |
| Tigriopus sp. | JQ688261.1 | South Korea | Tigriopus sp.-JQ688261.1-South Korea | 10.1071/IS17002 |

| | | | | |
|---|---|---|---|---|
| Tigriopus sp. | JQ688262.1 | South Korea | Tigriopus sp.-JQ688262.1-South Korea | 10.1071/IS17002 |
| Tigriopus sp. | JQ688263.1 | South Korea | Tigriopus sp.-JQ688263.1-South Korea | 10.1071/IS17002 |
| Tigriopus sp. | JQ688264.1 | South Korea | Tigriopus sp.-JQ688264.1-South Korea | 10.1071/IS17002 |
| Tigriopus sp. | JQ688265.1 | South Korea | Tigriopus sp.-JQ688265.1-South Korea | 10.1071/IS17002 |
| Tigriopus sp. | JQ688266.1 | South Korea | Tigriopus sp.-JQ688266.1-South Korea | 10.1071/IS17002 |
| Tigriopus sp. | JQ688267.1 | South Korea | Tigriopus sp.-JQ688267.1-South Korea | 10.1071/IS17002 |
| Tigriopus sp. | JQ688268.1 | South Korea | Tigriopus sp.-JQ688268.1-South Korea | 10.1071/IS17002 |
| Tigriopus sp. | JQ688269.1 | South Korea | Tigriopus sp.-JQ688269.1-South Korea | 10.1071/IS17002 |
| Tigriopus sp. | JQ688270.1 | South Korea | Tigriopus sp.-JQ688270.1-South Korea | 10.1071/IS17002 |
| Tigriopus sp. | JQ688271.1 | South Korea | Tigriopus sp.-JQ688271.1-South Korea | 10.1071/IS17002 |
| Tigriopus sp. | JQ688272.1 | South Korea | Tigriopus sp.-JQ688272.1-South Korea | 10.1071/IS17002 |
| Tigriopus sp. | JQ688273.1 | South Korea | Tigriopus sp.-JQ688273.1-South Korea | 10.1071/IS17002 |
| OUT-Acartia tonsa | MK319927.1 | | OUT-Acartia tonsa-MK319927.1 | 10.1007/s12526-020-01043-1 |
| OUT-Caligus clemensi | AM235887.1 | | OUT-Caligus clemensi-AM235887.1 | 10.3354/dao073141 |
| OUT-Lepeophtheirus salmonis | AY386274.1 | | OUT-Lepeophtheirus salmonis-AY386274.1 | Unpublished |
